# Human Cytomegalovirus uses a Host Stress Response to Balance the Elongation of Saturated/Monounsaturated and Polyunsaturated Very Long Chain Fatty Acids

**DOI:** 10.1101/2021.01.16.426974

**Authors:** Yuecheng Xi, Lena Lindenmayer, Ian Kline, Jens von Einem, John G. Purdy

**Affiliations:** Department of Immunobiology, University of Arizona, Tucson, Arizona, USA; Institute of Virology, Ulm University Medical Center, Ulm, Germany; BIO5 Institute, University of Arizona, Tucson, Arizona, USA; Cancer Biology Interdisciplinary Program, University of Arizona, Tucson, Arizona, USA

## Abstract

Stress and virus infection are known to regulate lipid metabolism in cells. Human cytomegalovirus infection induces fatty acid (FA) elongation and increases the cellular abundance of lipids with very long-chain FA tails (VLCFAs). While reprogramming of metabolism can be stress-related, the role of stress in HCMV reprogramming of lipid metabolism is poorly understood. In this study, we engineered cells to knockout PKR-like ER kinase (PERK) in the ER stress pathway and measured lipid changes using lipidomics to determine if PERK is needed for lipid changes associated with HCMV infection. We found that in HCMV-infected cells, PERK promotes the increase in the levels of phospholipids with saturated FA (SFA) and monounsaturated FA (MUFA) VLCFAs tails. Consistent with the SFA/MUFA lipidome changes, PERK enhances the protein levels of FA elongase 7 (ELOVL7), which elongates SFA and MUFA VLCFAs. Additionally, we found that increases in the elongation of polyunsaturated fatty acids (PUFAs) associated with HCMV infection was independent of PERK and that lipids with PUFA tails accumulated in HCMV-infected PERK knockout cells. Consistent with the PUFA lipidome changes, the protein levels of ELOVL5, which elongates PUFAs, are increased by HCMV infection through a PERK-independent mechanism. These observations show that PERK differentially regulates ELOVL7 and ELOVL5, creating a balance between the synthesis of lipids with SFA/MUFA tails and PUFA tails. Additionally, we found that PERK was necessary for virus replication and the infectivity of released viral progeny. Overall, our findings indicate that PERK—and more broadly, ER stress—may be necessary for membrane biogenesis needed to generate infectious HCMV virions.

**IMPORTANCE:** HCMV is a common herpesvirus that establishes lifelong persistent infections. While infection is asymptomatic in most people, HCMV causes life-threatening illnesses in immunocompromised people, including transplant recipients and cancer patients. Additionally, HCMV infection is a leading cause of congenital disabilities. HCMV replication relies on lipid synthesis. Here, we demonstrated that the ER stress mediator, PERK, controls fatty acid (FA) elongation and cellular abundance of several types of lipids following HCMV infection. Specifically, we found that PERK promotes FA elongase 7 synthesis of lipids with saturated/monounsaturated very long-chain FA tails which are important for building the viral membrane of infectious HCMV virions. Overall, our study shows that PERK is an essential host factor that supports HCMV replication and promotes lipidome changes caused by HCMV infection.

## INTRODUCTION

Regulation of metabolism in response to acute and chronic external or internal stressors allows cells to adapt to stressful situations, including viral infections (1–6). Human cytomegalovirus (HCMV) reprograms metabolism, in part by inducing stress responses, to create a metabolic state that supports virus replication (7, 8). The endoplasmic reticulum (ER) has roles in both lipid metabolism and stress. The integrated stress response, lipid synthesis, and protein translation and folding take place in the ER. Disruption of ER homeostasis triggers a stress response that can include activating PKR-like ER kinase (PERK, also known as eukaryotic translation initiation factor 2α kinase 3, EIF2AK3). PERK reduces ER-stress related to the accumulation of misfolded proteins by inactivating EIF2α through phosphorylation to decrease translation. PERK has an emerging role in lipid metabolism as well. PERK-mediated lipogenesis supports the proper development of lipid-generating tissues, such as adipocytes and mammary glands (9). In the context of viral infection, PERK is needed for HCMV-induced lipid synthesis (7, 10). However, the effects of PERK on specific classes or types of lipids in HCMV-infected cells remains unknown.

HCMV infection induces several lipogenic enzymes, including those involved in fatty acid (FA) synthesis and elongation (8, 10–14), which are required for HCMV replication (10, 12–14). FAs have several functions as free fatty acids or as hydrophobic tails in lipids, including establishing and maintaining membrane integrity. The function of FAs depends on their length and the number and placement of double bonds in the hydrocarbon chain. HCMV infection promotes the synthesis of lipids with very long-chain fatty acid (VLCFA) tails containing 24 or more carbons with no double bonds (i.e., saturated fatty acids, SFAs) or one double bond (i.e., monounsaturated fatty acids; MUFAs) (13, 14). In HCMV infected cells, SFAs are made by fatty acid elongase 7 (ELOVL7) (13). Humans encode seven ELOVLs that elongate FAs based on the chain length and double bond content (15–19). The expression of ELOVLs is regulated by sterol regulatory-element binding proteins (SREBPs), liver X receptor α (LXRα), and peroxisome proliferation-activated receptor α (PPARα) (20). However, little is known concerning mechanisms involved in the differential expression of ELOVLs (i.e., how a cell may regulate ELOVL7 differently than ELOVL5).

We recently demonstrated that HCMV UL37x1 protein (pUL37x1, also known as viral inhibitor of apoptosis, vMIA) helps to support HCMV enhancement of SFA elongation and synthesis of phospholipids (PLs) with VLCFA tails (PL-VLCFAs) (8). Additionally, pUL37x1 helps to increase PERK protein levels following infection (8). In this study, we investigated the role of PERK in the biogenesis of different lipids classes in response to HCMV infection. We engineered PERK knockout (KO) cells using CRISPR/Cas9 and performed several lipidomic analyses to define lipids regulated by PERK following HCMV infection. We expanded on our previous PL lipidomic work by including the analysis of diglycerides (DGs) and triglycerides (TGs). HCMV infection increases the relative levels of most DGs and TGs with SFA/MUFA VLCFA tails. We found that PERK affects the levels of PLs, DGs, and TGs, specifically in the double bond content of their tails. Our findings reveal that PERK balances the ratio of lipids with SFA/MUFA VLCFA tails and lipids with PUFA tails. Moreover, we show that in HCMV-infected cells PERK differentially regulates FA elongases (ELOVLs) involved in SFA/MUFA and PUFA elongation. Our study provides a new perspective on how HCMV remodels the host lipidome and how cellular stress regulates FA elongation and balances SFAs/MUFAs and PUFAs.

## RESULTS

### HCMV infection promotes PERK to support virus replication

HCMV infection with the lab-adapted AD169 strain increases PERK protein levels in primary human fibroblasts under fully confluent, serum-free conditions used in our metabolism studies (8, 13). We examined if infection with the low-passaged, more clinically-relevant TB40/E strain would also enhance PERK protein levels in the same cell culture conditions. PERK protein levels in mock-infected and TB40/E-infected cells were measured following infection at a multiplicity of infection (MOI) of 1 infectious unit per cell. PERK protein levels increased as early as 4 hours post-infection (hpi) and remained elevated through 96 hpi, with the highest levels observed at 24-72 hpi (Fig. 1A-B). Next, we examined if PERK is activated in HCMV-infected cells by measuring ATF4 protein levels, a transcription factor whose expression is increased by PERK activity. ATF4 levels were enhanced in HCMV-infected cells relative to uninfected cells at 48-120 hpi (Fig.1A and 1C). ATF4 expression was highest at 72 hpi with a 5.5-fold higher level in HCMV-infected cells than uninfected cells. Our findings that HCMV infection enhances PERK protein levels and activity is consistent with others that reached the same conclusion using other HCMV strains or cell types (5, 7, 8, 21, 22).

**Figure 1.**
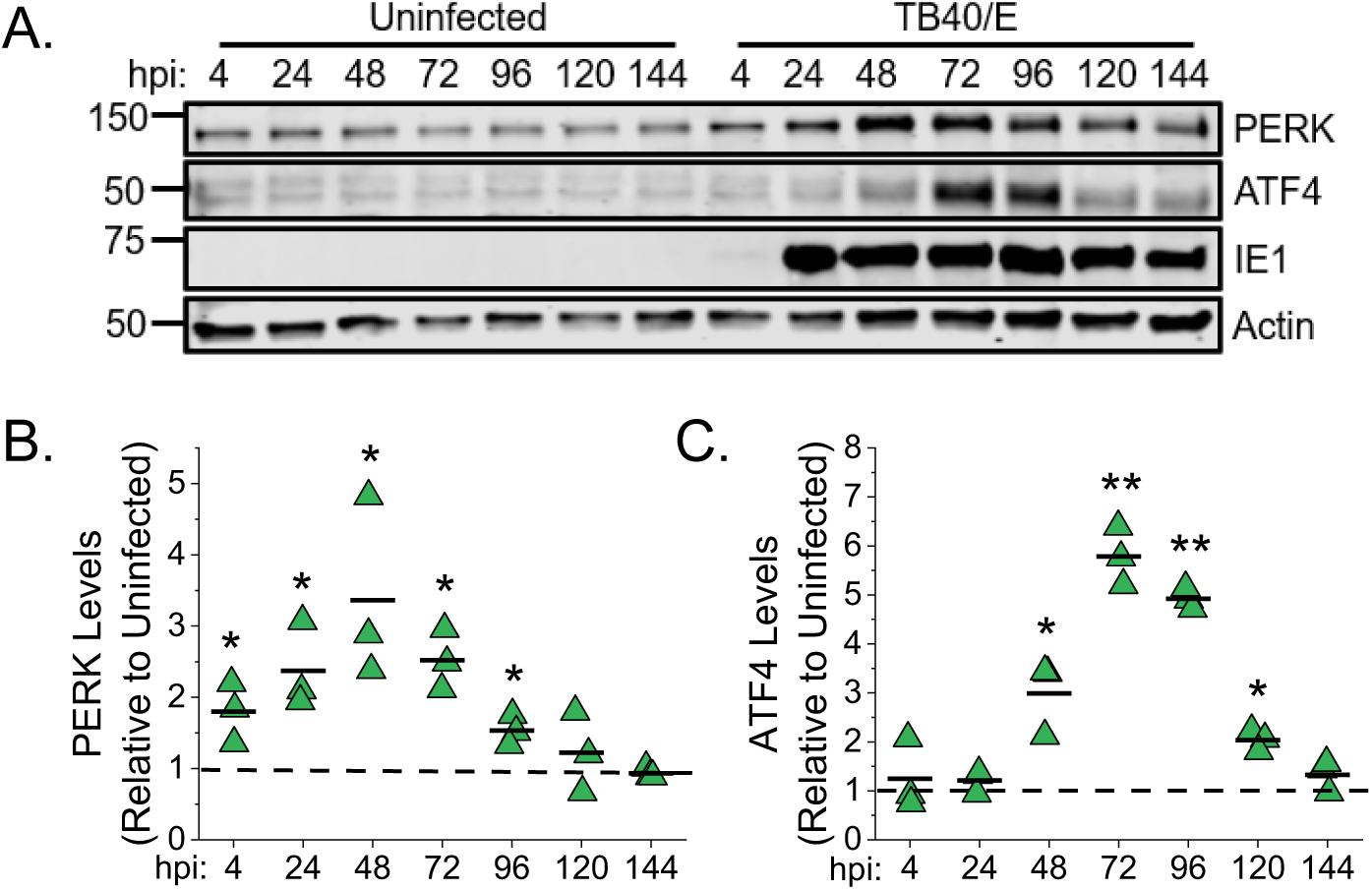
HCMV infection upregulates PERK and ATF4 protein levels. **(A)** Western blot analysis of PERK and ATF4 protein levels in uninfected and HCMV TB40/E-infected fibroblast cells at a multiplicity of infection (MOI) of 1 infectious unit per cell. **(B-C)** Quantification of PERK and ATF4 protein levels following normalization to the actin level. The value for PERK or ATF4 at each time point is shown relative to the level in uninfected cells at the same time point. A dashed line represents the level in uninfected cells and each data point from three independent experiments is shown with the mean represented as a bar. *p<0.05, **p<0.01, pair-sample t test. N=3.

To investigate possible functions of PERK in HCMV-infected cells, we generated PERK-knockout (KO) clones using CRISPR/Cas9 and two independent gRNAs (Fig. S1A-C). Each PERK-KO clone was generated by single-cell cloning using human foreskin fibroblast (HFF) cells exogenously expressing human telomerase. CRISPR/Cas9 gene editing was identified using Indel sequencing. Clone 1 (PERK-KO-c1) has a 300 base pairs (bp) deletion in exon one that removed the start codon for PERK translation (Fig. S1A-B). Clone 2 (PERK-KO-c2) contains a four base pair deletion after the PERK translation start site that introduces a frameshift leading to a premature stop codon. The loss of PERK protein was confirmed using Western blot (Fig. S1C). We generated CRISPR/Cas9-expressing control cells that contain a non-targeting (NT) gRNA sequence that lacks the ability to target any human or HCMV gene. The NT cells were single-cell cloned in parallel with the PERK-KO cells. Individual NT clones were pooled to limit off-target effects from CRISPR/Cas9 cloning.

The loss of PERK in the KO cells was further confirmed in HCMV-infected cells using Western blot. TB40/E-infected PERK-KO cells failed to express PERK protein (Fig. 2A). Next, we evaluated PERK activity in the KO cells by measuring the levels of ATF4 in TB40/E-infected cells. At 72 and 96 hpi, the levels of ATF4 protein were lower in HCMV-infected PERK-KO cells than in infected NT cells (Fig. 2A-B). The decrease in ATF4 protein levels confirms the loss of PERK activity in the KO cells.

**Figure 2.**
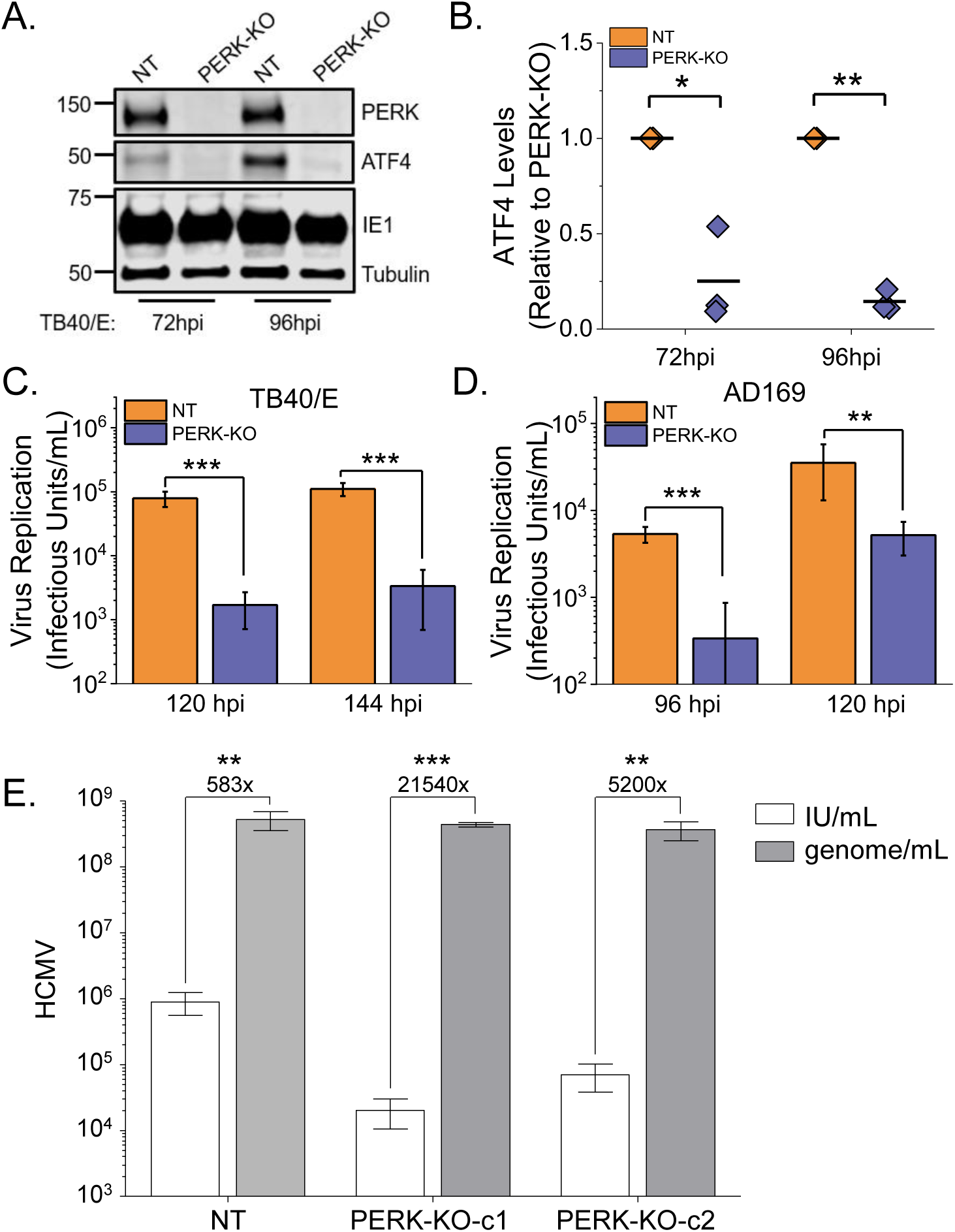
PERK enhances HCMV replication. **(A)** Western analysis of PERK and ATF4 protein levels in PERK-KO-c1 and NT cells at infected MOI of 1. **(B)** Quantification of ATF4 protein levels following normalization to tubulin protein levels. The value for ATF4 is shown relative to the level in TB40/E-infected NT cells. Each data point from three independent experiments is shown with the mean represented as a bar. **(C)** PERK-KO-c1 and NT cells were infected with TB40/E at a MOI of 1. The number of infectious intracellular and extracellular virus was measured at 120 hpi and 144 hpi by TCID50. **(D)** Release of infectious virus particles was assayed by infecting PERK-KO-c1 and NT cells with AD169 at a MOI of 1. At 96 hpi and 120 hpi, infectious extracellular virus released into the growth medium was measured by TCID50. **(E)** Virus yields in cell free supernatants were determined at 144 hpi following an infection at MOI of 3 (white bars). In the same supernatants, the number of genome-containing particles released by cells was measured by qPCR (gray bars). The genome-to-infectious unit (IU) ratio was determined, and the values are listed above the bars. *p<0.05, **p<0.01, ***p<0.001. One-sample t test for (B) and two-sample t test for (C-E). N=3.

The KO cells were used to determine if PERK is required for HCMV replication by infecting PERK-KO and NT cells with TB40/E at a MOI of 1. Since TB40/E infectious virus is both cell-associated and extracellularly released, we measured total virus production (i.e., cell-associated and released virions). At 120 and 144 hpi, infectious viral progeny was measured using tissue culture infectious dose 50 (TCID_50_). TB40/E virus progeny production was >10-fold lower in PERK-KO cells relative to NT control cells (Fig. 2C). We further defined the impact of PERK on HCMV infection by measuring virus cell-to-cell spread using a focus expansion assay. In this assay, cells were infected with 100 infectious HCMV virions, and at 24 hpi, the cells were overlaid with methylcellulose to limit virus spread from released particles. At 9 days post-infection (dpi), HCMV infected cells were visualized using immunofluorescence microscopy after staining for IE1, and the number of infected cells per plaque was determined. NT control cells had a mean of 38 HCMV-infected cells per plaque (Fig. S1D). In contrast, HCMV formed smaller foci with a mean of 21 infected cells per plaque in PERK-KO cells.

Next, we addressed the possibility that PERK may support the release of infectious progeny. We first addressed this possibility using the AD169 strain that is more efficiently released by infected fibroblast cells than TB40/E. We used TCID_50_ to measure the amount of infectious virus released at 96 and 120 hpi by infecting NT and PERK-KO cells using a MOI of 1. PERK-KO-c1 cells released infectious AD169 virus progeny 10-fold lower than NT cells (Fig. 2D). Release of AD169 progeny was also lower in PERK-KO-c2 cells relative to NT cells (Fig. S1E). We further confirmed these findings using cells infected at a MOI of 3 with TB40/E to increase the amount of released virus. At 144 hpi, PERK-KO cells infected at a MOI of 3 released >10-fold fewer infectious virus than NT cells (Fig. 2E, white bars), consistent with our findings using AD169.

Our observations suggest that PERK-KO cells release fewer infectious virus particles or that the released virus particles are less infectious than the virus particles made by NT cells. First, we examined the possibility that PERK supports HCMV release by measuring the total number of genome-containing viral particles released into the growth medium. At 144 hpi, the supernatant from PERK-KO and NT cells was collected, and the number of genomes present was determined by quantitative PCR. The number of genome-containing virus particles released by PERK-KO cells was similar to NT cells (Fig. 2E, gray bars). This finding indicates that PERK is not required for the production and release of virus particles. Next, we determined if the virus particles released by PERK-KO cells are less infectious than those released by NT cells to assess if PERK affects HCMV infectivity. Therefore, we calculated the ratio of genome-containing particles to infectious units (i.e., genome-to-IU ratio). The genome-to-IU ratio was around 580 in the control cells (Fig. 2E). In the PERK-KO cells, the genome-to-IU ratio was between 5,200 to 21,540. A higher ratio represents reduced particle infectivity. Since the genome-to-IU ratio was 8- to 36-fold greater in the PERK-KO than control cells, our findings indicate that PERK-KO cells release virus particles with reduced infectivity.

We conclude that PERK is necessary for efficient HCMV replication and the infectivity of released viral progeny. Moreover, our observations using CRISPR/Cas9 engineered PERK-KO cells are consistent with those observed in shRNA PERK knockdown cells (7), providing support that our observed phenotypes are due to the loss of PERK activity and not due to our genetic engineering approach.

### HCMV infection elevates the levels of diglycerides (DG) and triglycerides (TG)

In viral and non-infectious diseases ER-stress, and specifically PERK, regulates lipid metabolism (7, 9, 23, 24). Prior to determining the role of PERK in lipid metabolism of infected cells, we further defined the effects of HCMV infection on lipids. HCMV infection promotes lipid synthesis and increases the intracellular concentration of several lipids classes (7, 8, 10, 13, 25, 26). We have previously shown that HCMV increases the concentration of phospholipids, particularly those with VLCFA tails (PL-VLCFAs) (8). However, the levels of diglycerides (DGs) and triglycerides (TGs), including those with VLCFA tails, have been studied less. DG lipids are comprised of two FA tails attached to a glycerol lipid backbone. DGs are intermediates in the synthesis of both phospholipids and TG lipids. TGs have three FA tails attached to a glycerol backbone allowing these molecules to store FAs that can be used when needed. We identified and quantitatively measured the relative levels of DGs and TGs in HCMV-infected and uninfected cells using liquid-chromatography high-resolution tandem mass spectrometry (LC-MS/MS). Since DGs and TGs are neutral lipids, sodiated and ammoniated adducts that formed during electrospray ionization were used to identify and quantitate their levels. First, the levels of DGs and TGs were visualized in HCMV-infected cells relative to mock-infected cells using a plot where each adduct form is represented by a dot (Fig. 3). In these dot plots the DG or TG levels in infected cells are on the y-axis, and the levels in uninfected cells are on the x-axis. If the level of a DG or TG is the same in HCMV-infected and uninfected cells, then its dot will fall along the dashed linear line shown in each dot plot. At 96 hpi, most DGs are above the line, demonstrating that their levels are greater in TB40/E-infected cells than uninfected cells (Fig. 3A). Since all DGs, except for one, were elevated in HCMV-infected cells at 96 hpi, we examined earlier time points to determine when DGs levels are altered by infection. At 48 hpi, approximately 35% of DGs were more abundant in infected cells than uninfected cells (Fig. S2A). At 72 hpi, approximately 75% of DGs were more abundant (≥2-fold) in HCMV-infected cells relative to uninfected cells (Fig. S2D). Next, we examined if AD169 infection alters the levels of DGs. At 72 hpi, the level of most DGs was greater in AD169-infected cells than in uninfected cells (Fig. S2G), similar to TB40/E-infected cells at 72 and 96 hpi. In summary, HCMV infection increases the cellular levels of DGs by 48 hpi, and their levels continue to rise through 96 hpi.

**Figure 3.**
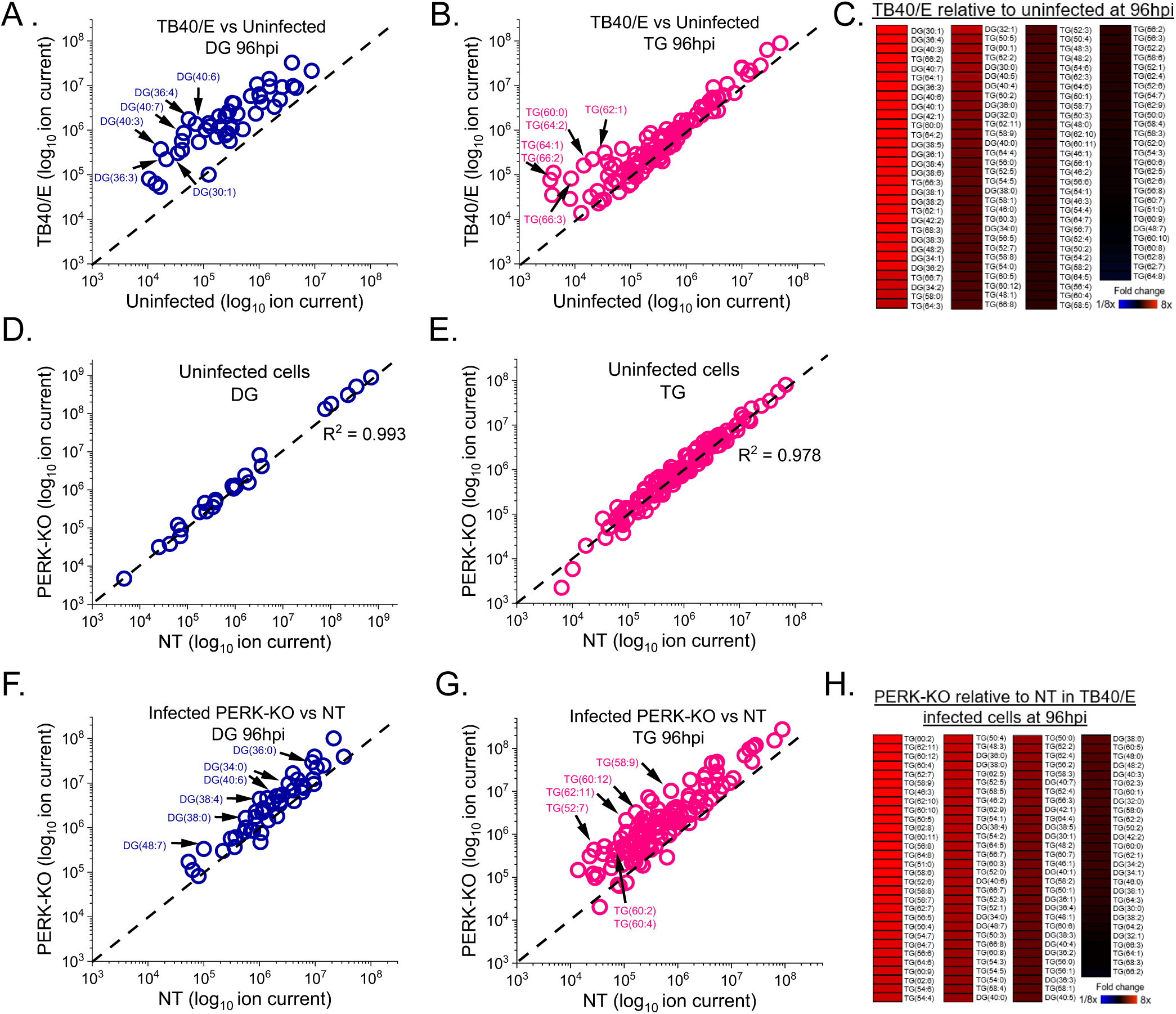
HCMV infection elevates diglycerides (DG) and triglycerides (TG), which are further increased by the loss of PERK. **(A)** Relative levels of DGs in TB40/E-infected cells and uninfected cells at 96 hpi. Sodiated and ammoniated adducts of DGs were measured by liquid chromatography high resolution tandem mass spectrometry (LC-MS/MS) using electrospray ionization (ESI) following normalization by cell number. Each dot in the plot represents the level of an adduct form of a DG lipid in TB40/E-infected cells relative to the its level in uninfected cells. The dash line represents a relative level of 1 (i.e., the level in infected cells is equal to the level in uninfected cells). **(B)** Relative levels of TGs in TB40/E-infected cells and uninfected cells at 96 hpi. TG data was analyzed and visualized using the same methods described in part A. **(C)** Changes in the relative levels of DGs and TGs in HCMV-infected cells and uninfected cells were quantified by averaging the relative levels of the sodiated and ammoniated adducts if both were measured. The averaged relative fold changes were log transformed and visualized as a heatmap. **(D-E)** Relative levels of DGs and TGs in uninfected PERK-KO and NT cells. **(F-H)** Relative levels of DGs and TGs in TB40/E-infected PERK-KO and NT cells at 96 hpi. DGs and TGs were analyzed and visualized as described in part A-C. All cells were infected at MOI of 3. N=3.

Since DGs form an intermediate step in the synthesis of TGs, the increased levels of DGs following HCMV infection may suggest that the levels of TGs could be altered by infection. At 96 hpi, ∼20% of TGs were more abundant (≥2-fold) in TB40/E-infected cells than uninfected cells (Fig. 3B). At 48 and 72 hpi, the levels of 14-20% of TGs were elevated in HCMV-infected cells relative to uninfected cells, respectively (Fig. S2B and S2E). Most of the TGs whose levels were increased by HCMV (e.g., TG(62:1), TG(62:2), and TG(64:2)) were elevated at all timepoints examined, indicating that they are increased by 48 hpi and remained elevated during virus replication. Several of the same TGs were more abundant in AD169-infected cells than uninfected cells at 72 hpi (Fig. S2H).

The dot plots visualize multiple adduct forms of DGs and TGs. To provide a more quantitative analysis when both the sodiated and ammoniated adduct forms were observed by LC-MS/MS, we averaged the relative fold change of their levels in infected to uninfected cells. The averaged relative fold changes were visualized using heatmaps. As the dot plots predicted, most DGs and TGs were >2-fold higher in TB40/E-infected cells than uninfected cells at 96 hpi (Fig. 3C). More specifically, the abundance of all DGs, except DG(48:7), and ∼20% of TGs was ≥2-fold higher in TB40/E-infected cells than in uninfected cells at 96 hpi. Similarly, DGs and TGs tended to be elevated in TB40/E-infected cells at 48 and 72 hpi, and AD169-infected cells at 72 hpi (Fig. S2C, F, and I).

### PERK-KO increases the levels of several DGs and TGs in HCMV-infected cells

We examined if PERK is required for the changes in DG and TG levels associated with HCMV infection by comparing the relative abundance of DGs and TGs in PERK-KO cells to NT cells. However, first, we determined if the loss of PERK affects DG and TG levels in uninfected cells in our experimental conditions, where cells are at full confluence and in serum-free growth medium. There was little to no difference in the levels of DGs and TGs between uninfected PERK-KO and NT cells (Fig. 3D-E). Next, we examined if the loss of PERK affects DG and TG levels in HCMV-infected cells. We infected PERK-KO and NT control cells with TB40/E at a MOI of 3. At 96 hpi, the levels of several DGs and TGs were greater in HCMV-infected PERK-KO cells than in infected NT cells (Fig. 3F-G). We found that 65% of DGs and 80% of TGs were ≥2-fold higher in HCMV-infected PERK-KO cells relative to infected NT cells (Fig. 3H). Likewise, most DGs and TGs were elevated in TB40/E-infected PERK-KO cells at 48 and 72 hpi (Fig. S3A-F). Next, we determined if PERK is required for changes in DG and TG levels following infection with AD169. Unlike TB40/E-infected cells, the levels of several DGs in AD169-infected PERK-KO cells were lower than in infected NT cells (Fig. S3G). However, AD169-infected PERK-KO cells had higher levels of TGs than infected NT cells, similar to the findings in TB40/E-infected cells (Fig. S3H-I).

Overall, the results show that HCMV infection increases the abundance of DG and TGs. The loss of PERK further promotes an increase in some lipids in TB40/E-infected cells. The observations presented in Figs. 3, S2, and S3 further suggest that the level of DGs may be HCMV strain-dependent depending on the presence of PERK.

### HCMV infection elevates the levels of DGs with PUFA tials

Our findings related to DGs described above, show that HCMV infection increases the relative abundance of most DG lipids. We sought to better understand if the DGs altered the most by HCMV infection were of a specific type. To address this, we examined the top six DGs whose levels were altered the greatest by TB40/E infection using the quantitative measurements visualized in our heatmaps. We further examined the top six DGs increased by infection as shown in the heatmap in Fig. 3C. At 96 hpi, the relative abundance of these six DGs were 10-40-fold greater in TB40/E-infected cells than uninfected cells (Fig. 4A). Most of these DGs contained polyunsaturated fatty acid tails (PUFAs) which have two or more double bonds (Fig. 4B). The PUFA tails had 3 to 6 double bonds and ranged in length from 20 to 22 carbons. While the relative abundance of these six DGs was greater in HCMV-infected cells compared to uninfected cells at 96 hpi, only half were increased at 72, suggesting that the DG levels in HCMV-infected cells are dynamic on a time scale of 1 day. In contrast to the TB40/E infection, the DGs increased the most by AD169 infection contained saturated fatty acid tails (SFA; i.e., those with no double bonds) or monounsaturated fatty acid tails (MUFA; i.e., those with one double bond) (Fig. S2G). However, it is noteworthy that in the AD169 experiments, fewer DGs with PUFA tails were measured as compared to the TB40/E experiments. These findings suggest that some DGs may be regulated in a strain specific manner. Nonetheless, we conclude that while that TB40/E infection increases the levels of most DGs, the levels of DG with PUFA tails are elevated the most by infection at late time points in virus replication.

**Figure 4.**
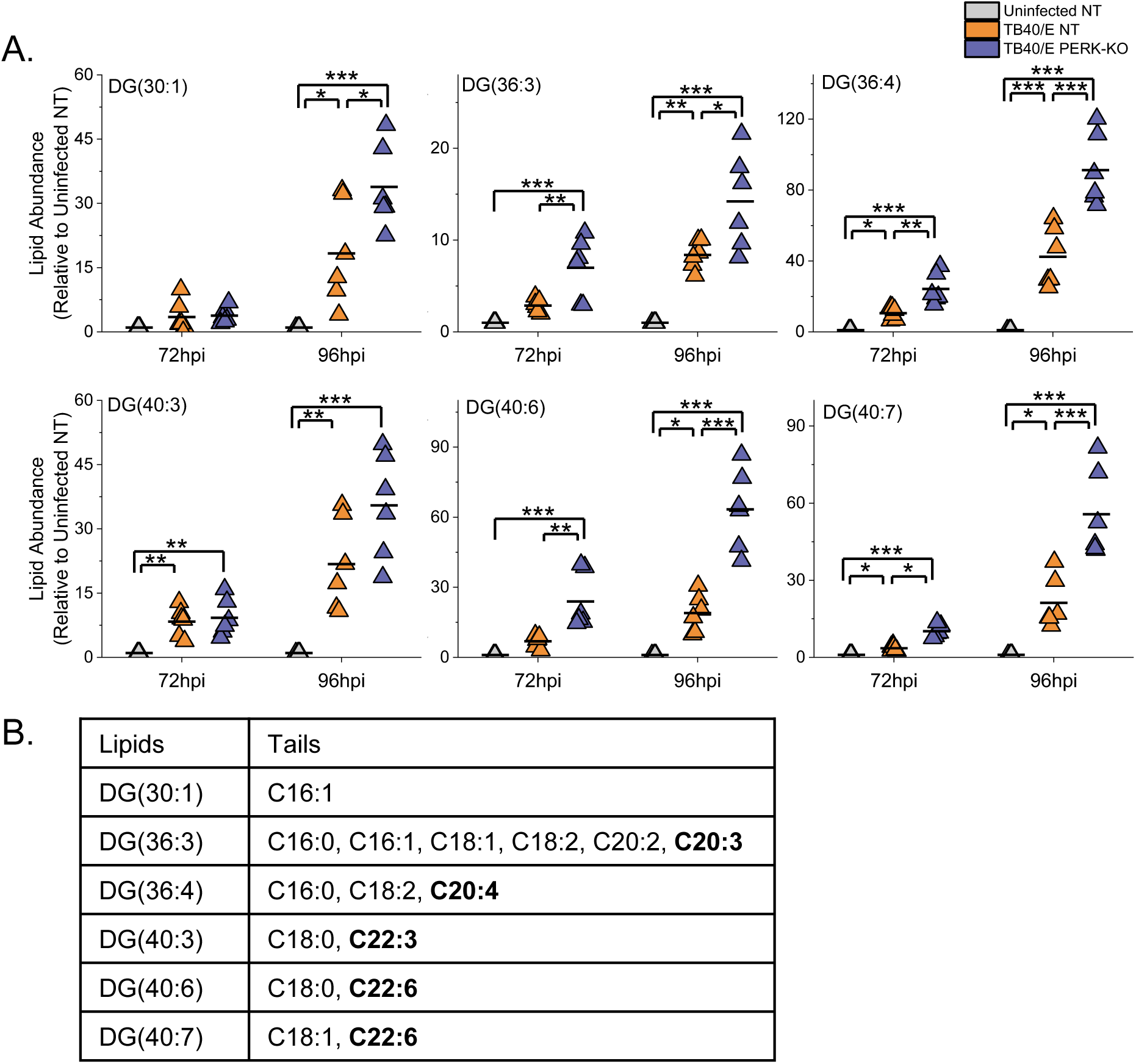
HCMV infection elevates DG lipids with long-chain and very long-chain polyunsaturated fatty acid tails. **(A)** The levels of DGs that were most prominently changed in TB40/E-infected cells relative to uninfected cells at 96 hpi. These are the top six elevated DGs shown in the heatmap of Fig. 3C. Three independent experiments were performed, each with duplicated samples for a total of six data points. Each data point is graphed relative to the levels observed in uninfected NT cells. For comparison, the relative levels at 72 hpi are also shown. *p<0.5, **p<0.01, ***p<0.001. One-way ANOVA, Turkey test. **(B)** The fatty acid tails for DGs shown in part A were identified by LC-MS/MS. Long-chain and very long-chain polyunsaturated tails (LCFA-PUFAs and VLCFA-PUFAs) are in bold text.

### The loss of PERK elevates the levels of DGs regardless of their tail composition

Since HCMV-infected cells have elevated levels of DGs with PUFA tails, we sought to determine if PERK influences the levels of DGs with PUFA tails during infection. Examination of the six DGs that are increased the most by HCMV infection (from Fig. 3C) showed further elevated levels of DGs in TB40/E-infected PERK-KO cells relative to infected NT cells and uninfected cells (Fig. 4A). Moreover, these DGs were more abundant in PERK-KO cells than infected control cells and uninfected cells at 72 hpi as well (Fig. 4A). As described above, most of these DGs have PUFA tails, suggesting that the loss of PERK leads to an accumulation of lipids with PUFA tails in HCMV-infected cells. Next, we determined if the loss of PERK would elevate other types of DGs, such as those with SFA or MUFA tails. We examined this possibility by identifying the tail composition of the six DGs that were increased the most in by the loss of PERK in HCMV-infected cells at 96 hpi (these are the top DGs shown in the heatmap in Fig. 3H). At 96 hpi, the relative abundance of these six DGs were 2- to 6-fold greater in infected PERK-KO cells than infected NT cells and 2.5- to 60-fold more abundant when compared to uninfected cells (Fig. 5A). Half of these DGs have PUFA tails, and the other half have SFA tails (Fig. 5B). Together, the findings shown in Figs. 4-5 indicate that the loss of PERK in HCMV-infected cells promotes the levels of DGs regardless of tail composition.

**Figure 5.**
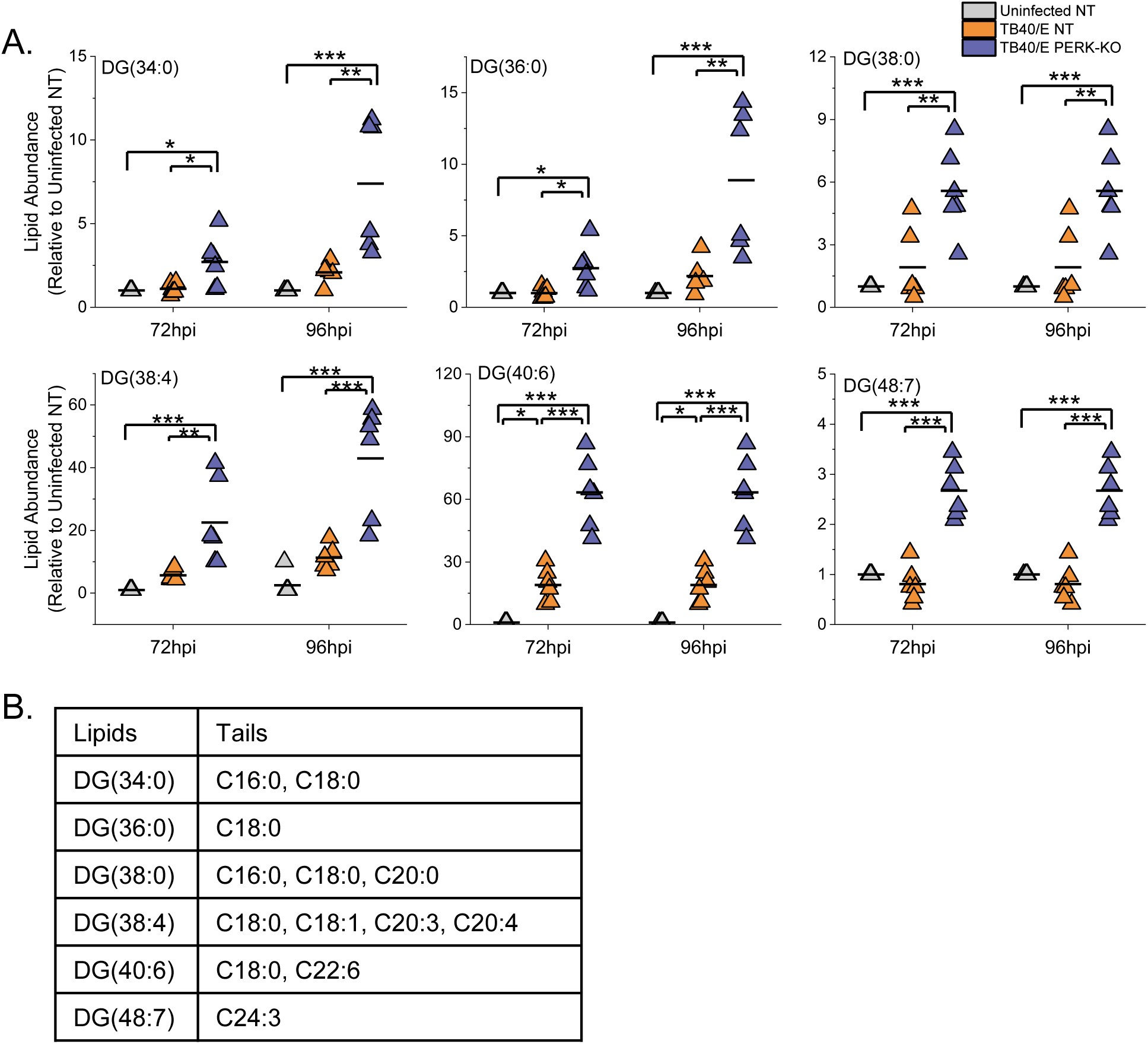
Loss of PERK leads to an accumulation of DGs regardless of tail compostion in HCMV-infected cells. **(A)** The levels of DGs that were most prominently changed in TB40/E-infected PERK-KO cells relative to infected NT cells at 96 hpi. These represent the top six elevated DGs shown in the heatmap of Fig. 3H. Each data point represents a sample from three independent experiments and is graphed relative to the levels observed in uninfected NT cells. For comparison, the relative levels at 72 hpi are also shown. *p<0.5, **p<0.01, ***p<0.001. One-way ANOVA, Turkey test. **(B)** The fatty acid tails for DGs shown in part A were identified by LC-MS/MS.

### HCMV infection increases TGs with SFA/MUFA VLCFA tails

Since HCMV infection increases the levels of most TGs, we further characterized the TGs with the greatest relative abundance between infected cells and uninfected cells at 96 hpi. We focused on the top six TGs shown in the heatmap in Fig. 3C. The relative abundance of each of these six TGs was 10- to 18-fold higher in HCMV-infected cells than in uninfected cells at 72 and 96 hpi (Fig. 6A). Each of these TGs had 60 or more total carbons in their tails. TGs have three FA tails, and several molecular forms with different combinations of tails may be present in each of these TGs. The tails in these TGs were identified using tandem MS/MS. At least four tails per TG were identified, demonstrating that each TG represents at least two molecular forms. Furthermore, each TG had at least two VLCFA tails with 24 or more carbons (≥C24) (Fig. 6B). Most of the VLCFA tails were saturated fatty acid (SFA; no double bond) or monounsaturated fatty acid (MUFA; a single double bond). In addition to the SFA and MUFA tails, TG(66:2) and TG(66:3) each had a tail with two double bonds. These results demonstrate that HCMV infection increases the levels of TGs with SFA/MUFA VLCFA tails.

**Figure 6.**
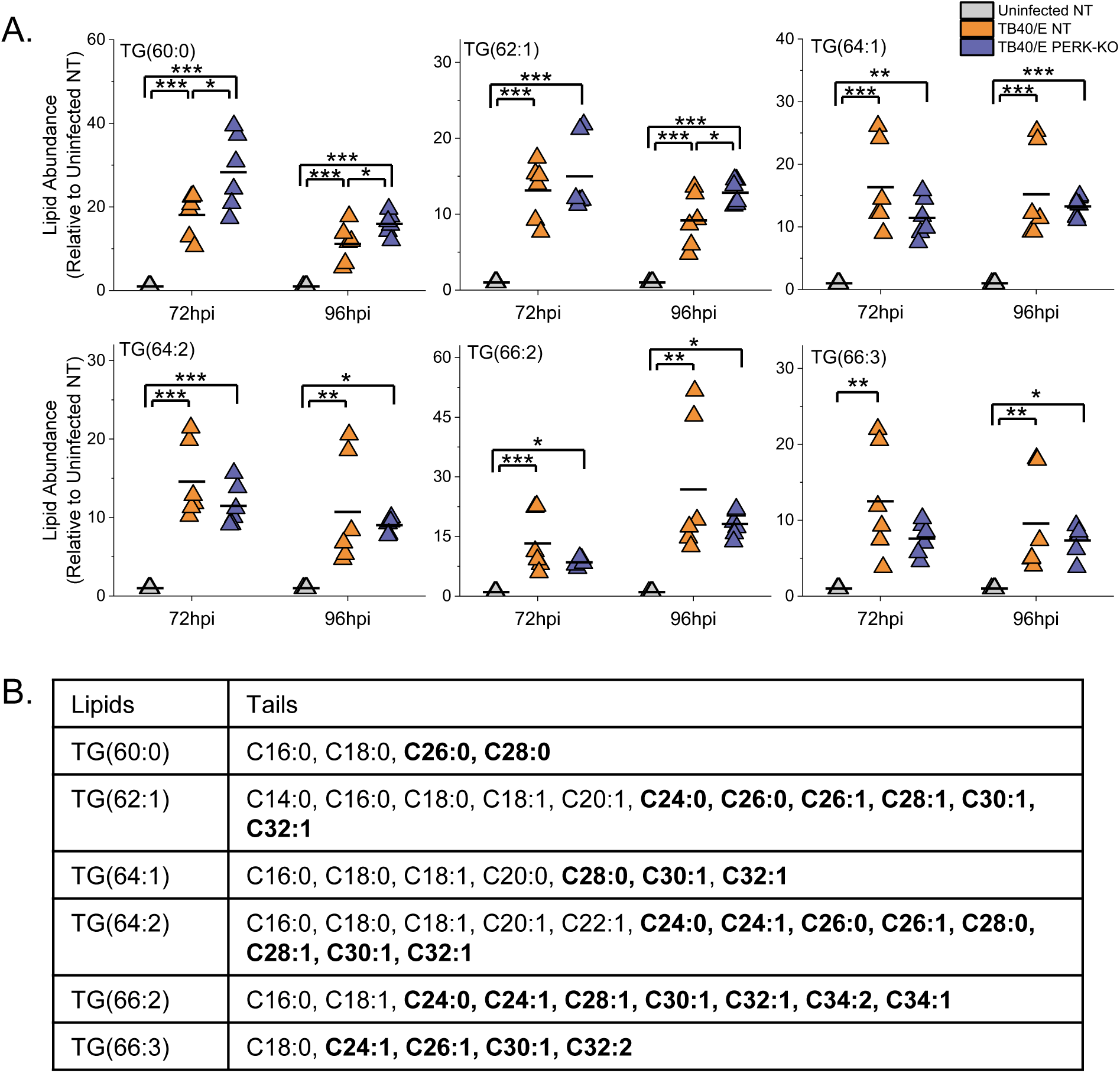
HCMV induces TGs with very long-chain saturated (SFA) or monounsaturated fatty acid (MUFA) tails independent of PERK. **(A)** The levels of TGs that were most prominently changed in TB40/E-infected cells relative to uninfected cells at 96 hpi. These are the top six elevated TGs shown in the heatmap of Fig. 3C. Three independent experiments were performed, each with duplicated samples for a total of six data points. Each data point is graphed relative to the levels observed in uninfected NT cells. For comparison, the relative levels at 72 hpi are also shown. *p<0.5, **p<0.01, ***p<0.001. One-way ANOVA, Turkey test. **(B)** The fatty acid tails for TGs shown in part A were identified by LC-MS/MS. SFA and MUFA VLCFA tails are in bold text.

### The loss of PERK elevates the levels TGs with PUFA tails in HCMV-infected cells

We continued to define the relationship between PERK and the tail composition by examining TGs. Of the six TGs increased the most by HCMV infection, four of the six had levels similar in TB40/E-infected PERK-KO and infected NT cells (Fig. 6A). At 72 and 96 hpi, the levels of TG(60:0) were greater in PERK-KO cells relative to NT cells (Fig. 6A). The same is true for TG(62:1) at 96 hpi. These results indicate that HCMV infection increases the levels TGs with SFA/MUFA VLCFA tails largely independent of PERK.

Next, we examined the TGs that were altered the most by the loss of PERK in HCMV-infected cells (i.e., the top six TGs in the heatmap shown in Fig. 3H). For each of these six TGs, their levels were 10- to 20-fold higher in TB40/E-infected PERK-KO cells relative to infected NT cells at 96 hpi (Fig. 7A). At 72 hpi, most of these TGs were also increased in infected PERK-KO cells relative to infected NT cells, but at a reduced level. Each of these TGs had at least four tails, demonstrating that all of them have multiple molecular forms (Fig. 7B). Five of the top six TGs contained multiple PUFA tails that were 20 to 26 carbons long (i.e., PUFA VLCFA tails). None of these TGs were higher in HCMV infected NT cells relative to uninfected NT cells, and one, TG(60:4), was slightly reduced by infection in NT cells at 72 hpi. These observations indicate that in the absence of PERK HCMV infection increases the levels of TGs with PUFA VLCFAs that are not altered by infection when PERK is present.

**Figure 7.**
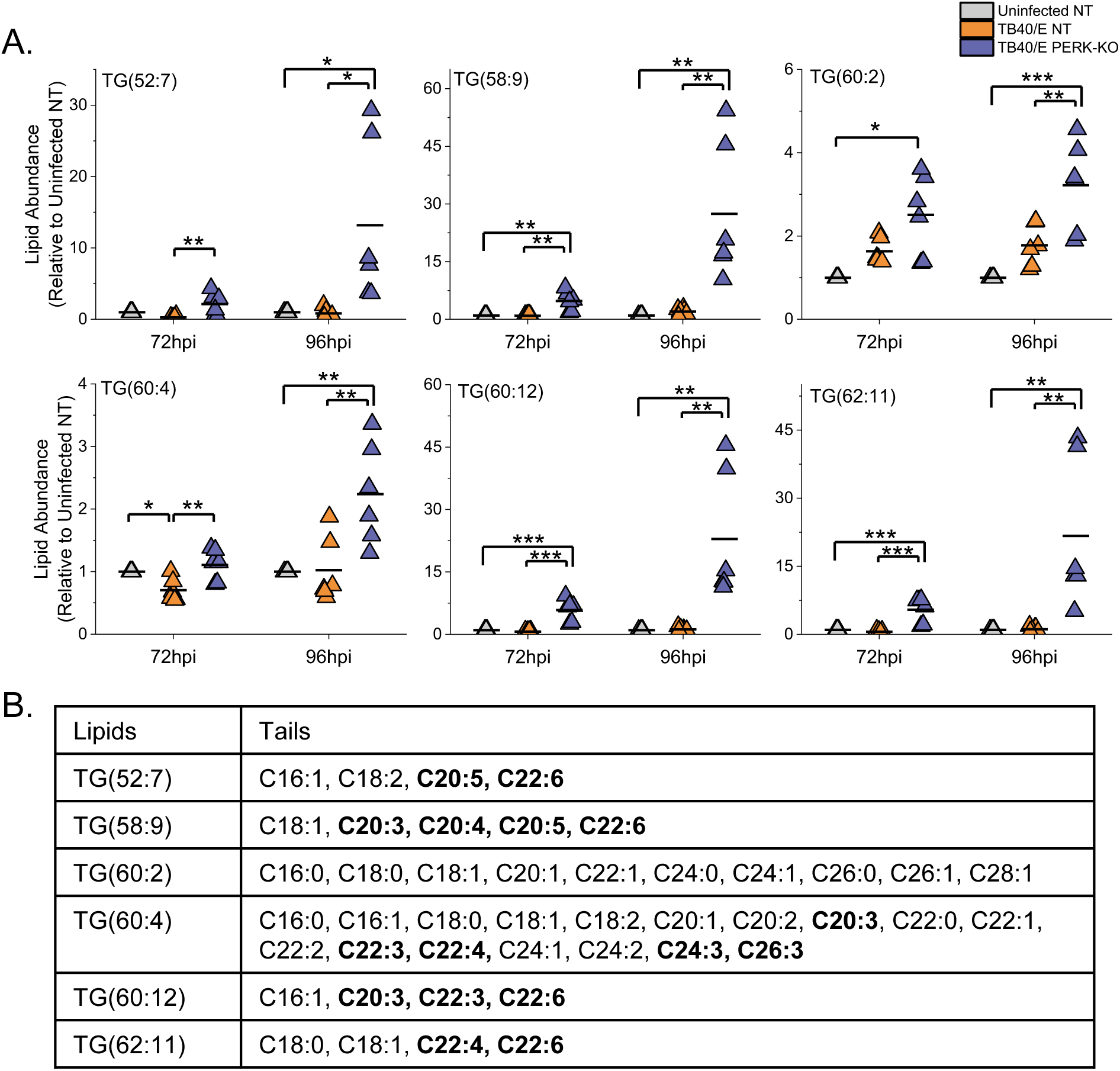
Loss of PERK leads to an accumulation of TGs with polyunsaturated fatty acid tails (PUFA) in HCMV-infected cells. **(A)** The levels of TGs that were most prominently changed in TB40/E-infected PERK-KO cells relative to infected NT cells at 96 hpi. These represent the top six elevated TGs shown in the heatmap of Fig. 3H. Each data point represents a sample from three independent experiments and is graphed relative to the levels observed in uninfected NT cells. For comparison, the relative levels at 72 hpi are also shown. *p<0.5, **p<0.01, ***p<0.001. One-way ANOVA, Turkey test. **(B)** The fatty acid tails for TGs shown in part A were identified by LC-MS/MS. PUFA tails are in bold text.

In summary, HCMV infection elevates the levels of TGs with SFA/MUFA VLFCAs independent of PERK (Fig. 6). Furthermore, in HCMV-infected cells the loss of PERK promotes the levels of TGs with PUFA VLCFAs. Together, our observations in Figs. 6-7 indicate that in HCMV-infected cells PERK provides a balance between TGs with SFA/MUFA VLCFA tails and those with PUFA VLCFA tails.

### HCMV infection increases the levels of phospholipids (PLs) with SFA/MUFA VLCFA tails

If PERK balances SFA/MUFA and PUFA tails during HCMV infection, we would expect that the tails of phospholipids (PLs) will also be dysregulated in HCMV-infected PERK-KO cells. We determined if PERK affects the abundance of phospholipids following HCMV infection by identifying and measuring the relative concentration of phospholipids using LC-MS/MS.

First, we determined if HCMV infection alters the levels of PLs by comparing the relative abundance of PLs in TB40/E-infected cells to uninfected cells at 96 hpi. At 96 hpi, 52% of the PLs were upregulated by ≥2-fold in HCMV-infected cells relative to uninfected cells (Fig. 8A-B). In contrast, the levels of only two lipids were downregulated by ≥2-fold infection (Fig. 8B). Since TB40/E-infection increases the levels of PLs at 96 hpi, we examined earlier timepoints to establish if TB40/E alters PLs levels at 48 and 72 hpi. At 48 hpi, the levels of ∼20% of PLs were ≥2-fold higher in HCMV-infected cells relative to uninfected cells (Fig. S4A-B). At 72 hpi, 34% of PLs were more abundant (≥2-fold) in infected cells than in uninfected cells (Fig. S4C-D). At 48 and 72 hpi, none of the PLs were reduced by HCMV infection (Fig. S4A-D). These findings show that the levels of several PLs are increased in cells by infection as early as 48 hpi, and continue to be elevated at 72 and 96 hpi. Next, we determined if AD169-infection alters PLs levels. At 72 hpi, the levels of several classes of PLs were greater in AD169-infected cells than in uninfected cells (Fig. S4E-F). In both AD169-infected and TB40/E-infected cells, many of the lipids that are elevated the most in infected cells are PLs with SFA/MUFA (Figs. 8A-B and S4). The observations reported here confirm our previous findings that infection with AD169 elevates the abundance of PLs with SFA/MUFA tails, particularly of PLs with SFA/MUFA VLCFAs with 24 or more carbons (8).

**Figure 8.**
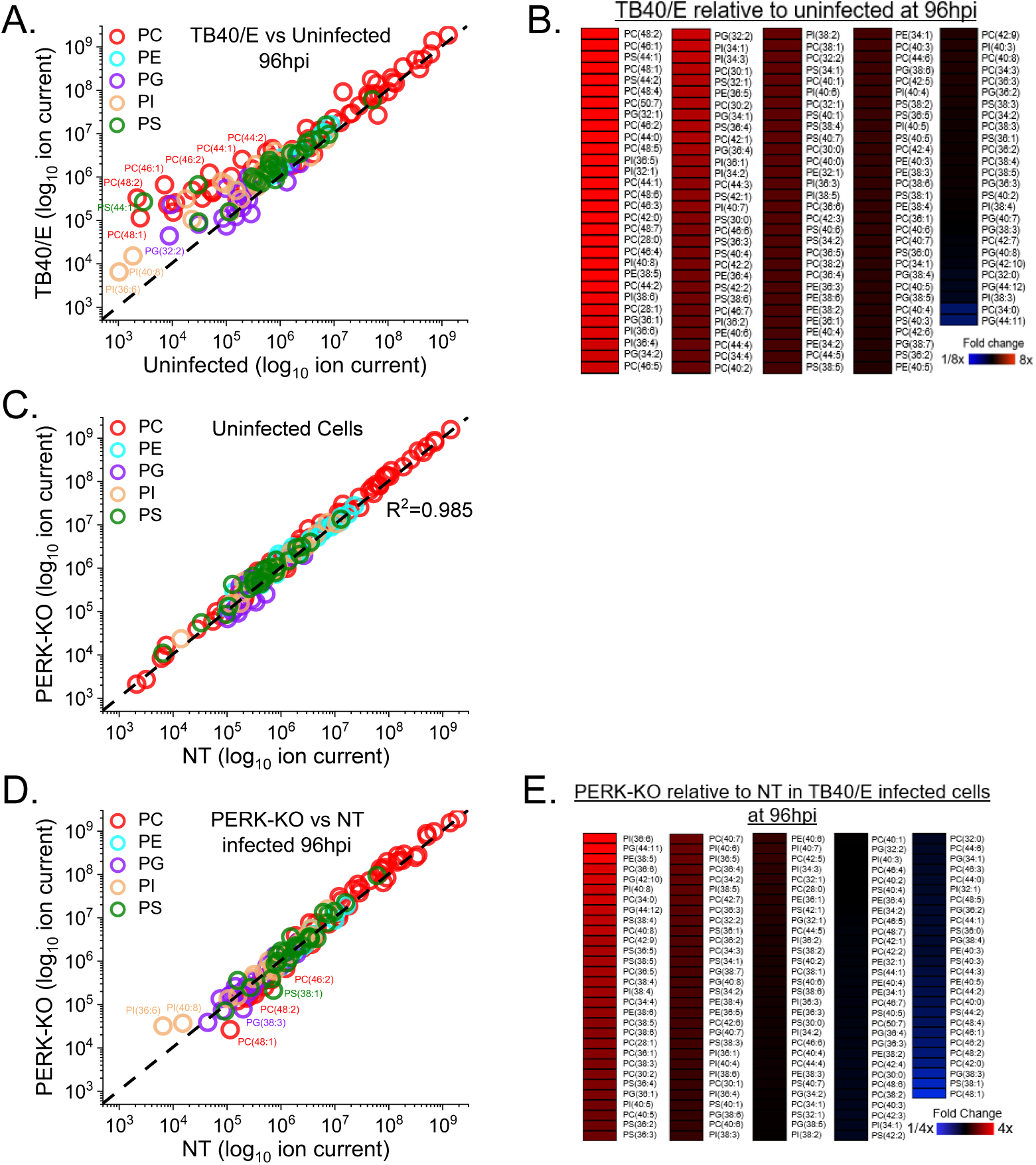
HCMV infection elevates phospholipids (PLs), which are reduced by the loss of PERK. **(A-B)** Relative levels of PLs in TB40/E-infected and uninfected cells at 96 hpi. PLs were measured by LC-MS/MS following normalization by cell number. Each dot in the plot represents the level of a PL in TB40/E-infected cells relative to the its level in uninfected cells. The dash line represents a relative level of 1 (e.g., the level in infected cells is equal to the level in uninfected cells). **(C)** Relative levels of PLs in uninfected PERK-KO and NT cells. **(D-E)** Relative levels of PLs in TB40/E-infected PERK-KO and NT cells at 96 hpi. Abbreviation: PC, phosphatidylcholine; PE, phosphatidylethanolamine; PG, phosphatidylglycerol; PI, phosphatidylinositol; and PS, phosphatidylserine. MOI=3. N=3.

We further characterized the PLs with the greatest relative abundance between infected cells and uninfected cells at 96 hpi. We focused on the top six PLs shown in the heatmap in Fig. 8B. The relative abundance of each of these six PLs was 10- to 150-fold higher in HCMV-infected cells than in uninfected cells at 72 and 96 hpi (Fig. 9A). Each of these PLs had 44 or more total carbons in their tails. These PLs had two tails, and several molecular forms with different tail combinations may be present in each. Half of these PLs had only two tails indicating the presence of a single molecular form. The others represent at least two molecular forms since we identified at least three tails per PL. Furthermore, each of these PLs contained a SFA or MUFA VLCFA tail with 26 or more carbons (Fig. 9B). These results further demonstrate that HCMV infection increases the levels of PLs with SFA/MUFA VLCFA tails.

**Figure 9.**
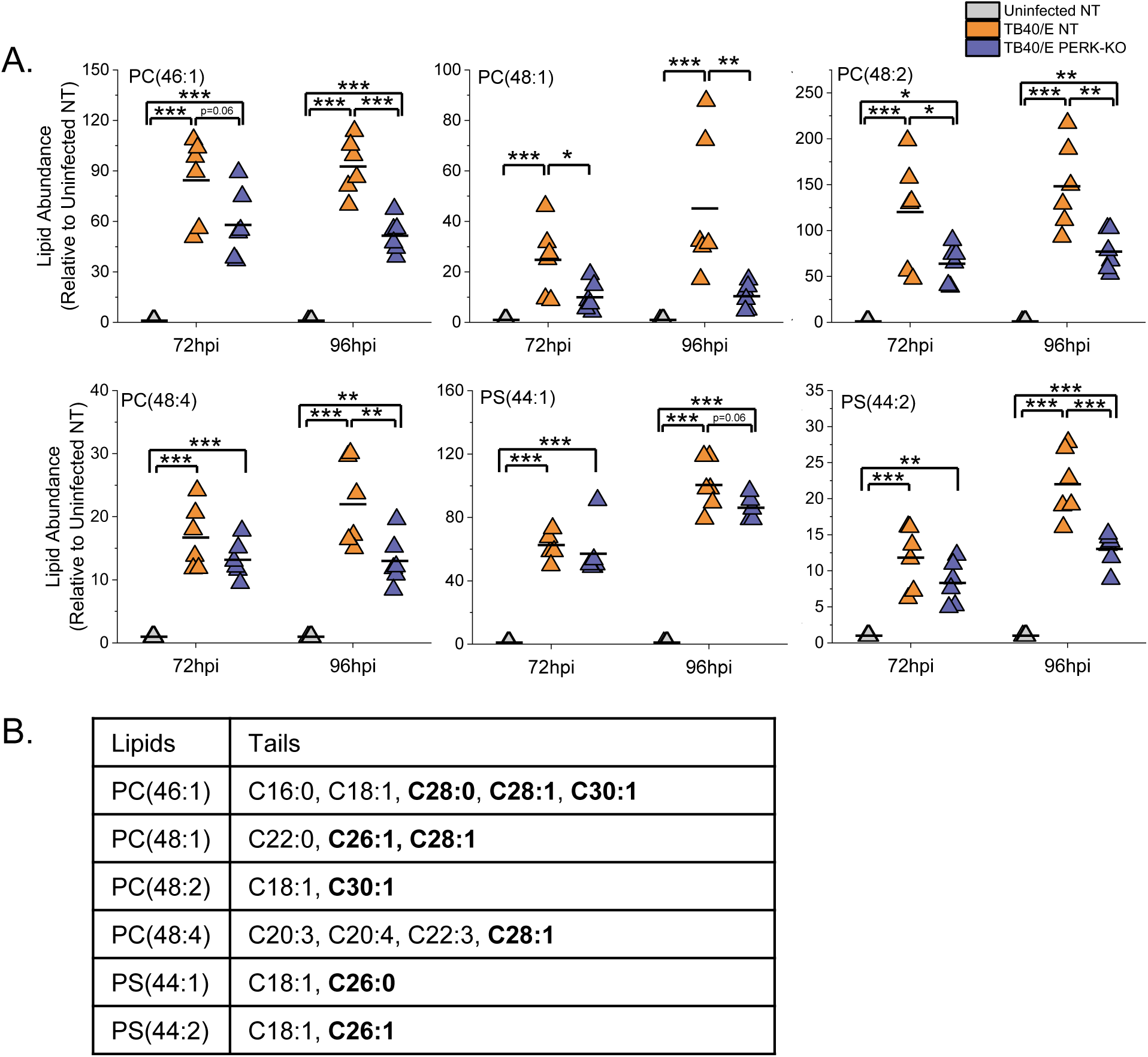
HCMV infection elevates PLs with SFA and MUFA VLCFA tails that is in part dependent on PERK. **(A)** Relative abundance of PLs that were most prominently changed in TB40/E-infected cells relative to uninfected cells at 96 hpi. These are the top six elevated PLs shown in the heatmap of Fig. 8B. Three independent experiments were performed, each with duplicated samples for a total of six data points. Each data point is graphed relative to the levels observed in uninfected NT cells. For comparison, the relative levels at 72 hpi are also shown. *p<0.5, **p<0.01, ***p<0.001. One-way ANOVA, Turkey test. **(B)** Tail composition for PLs shown in part A. SFA and MUFA VLCFA tails are in bold text.

### The loss of PERK alters the levels of PLs in HCMV-infected cells

Since the loss of PERK affects the levels of DGs and TGs, we considered if PERK also affects the levels of PLs in HCMV-infected cells. First, we compared the abundance of PLs in uninfected PERK-KO and NT cells to determine if PERK regulates the levels of PLs independent of infection. The levels of PLs in PERK-KO cells are equal to the levels in NT cells, demonstrating that the depletion of PERK has no significant effect on the abundance of PLs in uninfected cells (Fig. 8C). This observation was similar to the DGs and TGs profile in uninfected PERK-KO and NT cells, providing further evidence to support that PERK does not regulate lipid metabolism in uninfected cells under the conditions used in our experiments.

Next, we examined if the loss of PERK affects PL levels in HCMV-infected cells. We infected PERK-KO and NT control cells with TB40/E at a MOI of 3. At 96 hpi, the levels of several PLs were greater in HCMV-infected PERK-KO cells than in infected NT cells (Fig. 8D-E). We found that ∼16% of PLs were ≥2-fold more abundant in HCMV-infected PERK-KO cells relative to infected NT cells (Fig. 8D-E). Many of the PLs elevated in TB40/E-infected PERK-KO cells at 96 hpi were also elevated at 72 hpi (Fig. S5A-B). Next, we determined if PERK was required for changes in PL levels following infection with AD169. Like TB40/E-infected PERK-KO cells, the levels of several PLs in AD169-infected PERK-KO cells were greater than in infected NT cells (Fig. S5C-D).

### The loss of PERK reduces the levels of PLs with SFA/MUFA VLCFA tails

Some PLs were reduced in HCMV-infected PERK-KO cells. Approximately 6-12% of PLs in TB40/E- and AD169-infected PERK-KO were reduced by ≥2-fold compared to infected NT cells (Figs. 8D-E and S5). The levels of several of these PLs are increased by HCMV infection in NT cells but reduced by the loss of PERK, suggesting that PERK supports HCMV-induced increases in PL abundance. To further investigate this possibility, we focused on the PLs that were elevated the most in TB40/E-infected cells relative to uninfected cells at 96 hpi (i.e., the lipids shown in Fig. 8B). Of the six PLs elevated the most by HCMV infection, most were reduced in infected PERK-KO cells relative to infected NT cells (Fig. 9A). For example, PC(48:2) was the PL with the highest level of increase in TB40/E-infected cells relative to uninfected cells at 48, 72, and 96 hpi (Figs. 8A-B, 9A, and S4A-D). PC(48:2) was also among the largest PL changes following AD169 infection (Fig. S4E-F). However, in TB40/E-infected or AD169-infected PERK-KO cells, the level of PC(48:2) was significantly reduced relative to infected NT cells (Figs. 8D-E, 9A, and S5). These findings suggest that PERK promotes the levels of PLs with SFA/MUFA VLCFA tails in HCMV-infected cells. While the levels of these lipids are lower in HCMV-infected PERK-KO relative to infected NT cells, their levels in infected PERK-KO cells are still higher than in uninfected cells (Fig. 9A). For example, PC(48:2) is >50-fold more abundant in TB40/E-infected PERK-KO cells than uninfected cells (Fig. 9A). This observation suggests that PERK is not required to induce changes in lipid levels following HCMV infection, but may be necessary to promote changes in the lipidome. Overall, we conclude that PERK helps promote the levels of PLs with SFA/MUFA VLCFA tails in HCMV-infected cells.

### The loss of PERK elevates the levels of PLs with PUFA tails

We sought to understand further the role of PERK in regulating PL levels in HCMV-infected cells by identifying the lipids that were increased the most by the loss of PERK in HCMV-infected cells (i.e., the top six PLs in the heatmap shown in Fig. 8E). For each of these six PLs, their levels were >2.5-fold higher in TB40/E-infected PERK-KO cells relative to infected NT cells at 96 hpi (Fig. 10A). At 72 hpi, most of these PLs were also increased in infected PERK-KO cells relative to infected NT cells. Relative to uninfected NT cells, these lipids were 3- to 45-fold more abundant in HCMV-infected PERK-KO cells. However, their levels were similar in HCMV-infected NT cells and uninfected NT cells, demonstrating that HCMV infection does not significantly alter their abundance when PERK is present (Fig. 10A). These PLs were PI, PG, PE, and PC lipids. We identified tails for four of these lipids but were unable to identify the tails for the PI lipids using our MS/MS approach. These PLs contained PUFA tails ranging in length from 20 to 22 carbons (Fig. 10B). Based on the observations shown in Fig. 10, we conclude that the loss of PERK promotes the levels of PLs with PUFA tails in HCMV-infected cells.

**Figure 10.**
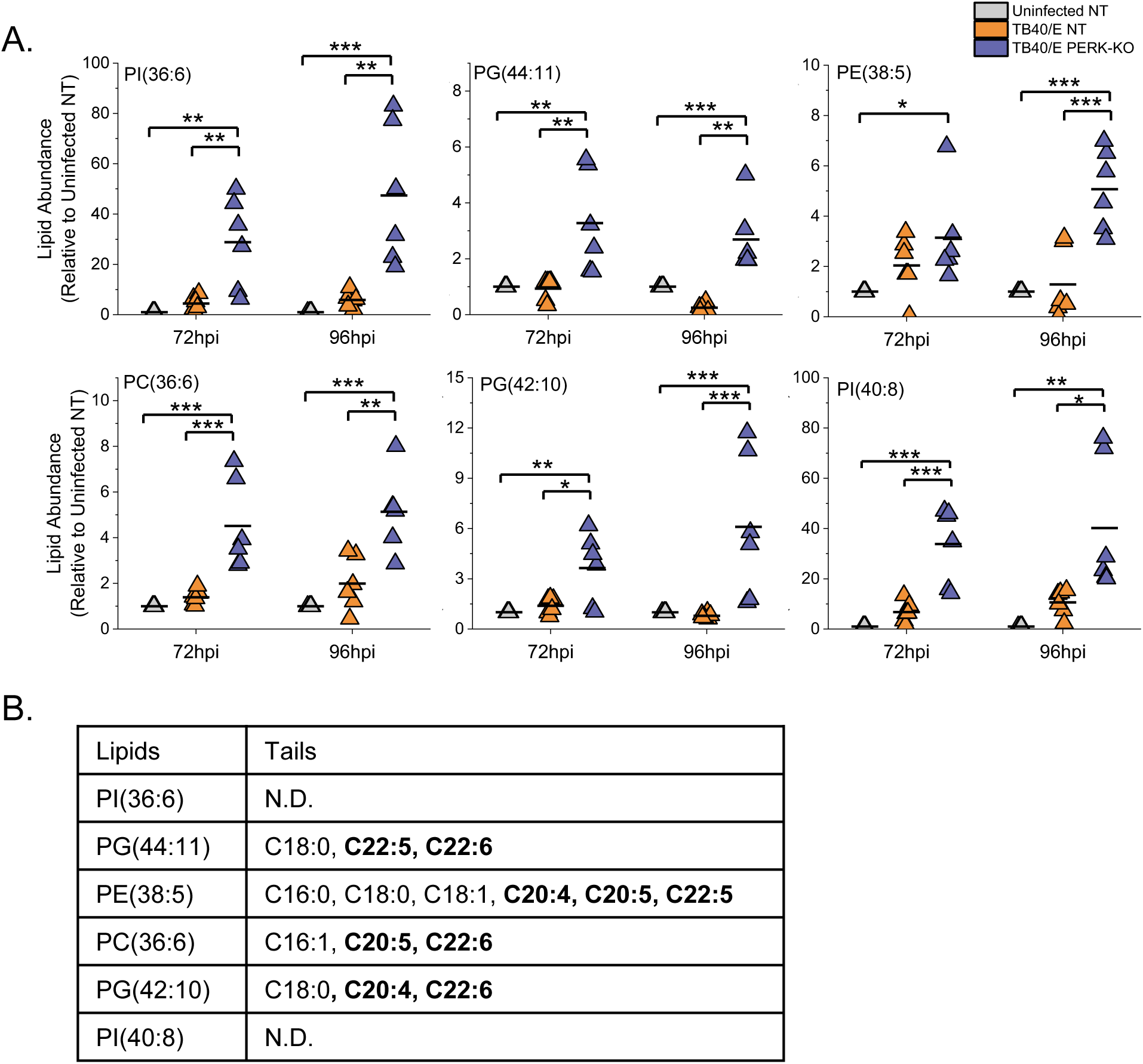
Loss of PERK leads to an accumulation of PLs with PUFA tails in HCMV-infected cells. **(A)** Relative abundance of PLs that were most prominently changed in TB40/E-infected PERK-KO cells relative to infected NT cells at 96 hpi. These are the top six elevated PLs shown in the heatmap of Fig. 8E. Each data point is graphed relative to the levels observed in uninfected NT cells. For comparison, the relative levels at 72 hpi are also shown. *p<0.5, **p<0.01, ***p<0.001. One-way ANOVA, Turkey test. **(B)** Tail composition for PLs shown in part A. PUFA LCFA and VLCFA tails are in bold text.

### PERK enhances ELOVL7 protein levels but not ELOVL5 levels

The lipid analysis demonstrated that PERK regulates a balance between SFA/MUFA VLCFA tails and PUFA tails in TGs and PLs in HCMV-infected cells. ELOVL7 elongates SFA and MUFA VLCFAs (Fig. 11A) (13, 15, 16, 27). We previously demonstrated that HCMV infection increases ELOVL7 gene expression (13). Protein levels of ELOVL7 were also upregulated by 17-20-fold in HCMV-infected HFF cells compared to uninfected at 72 and 96 hpi, respectively (Fig. 11B-C). ELOVL7 is also required for HCMV replication (13). In addition to ELOVL7, HCMV replication relies on ELOVL5 (13). ELOVL5 elongates PUFA VLCFAs (Fig. 11A) (15, 19). HCMV infection also increases ELOVL5 gene expression (13) and protein levels by 2.5-fold at 72 and 96 hpi (Fig. 11D-E). We hypothesized that during HCMV replication, PERK balances SFAs/MUFAs and PUFAs by regulating ELOVL7 and ELOVL5 protein levels. To test our hypothesis, we infected PERK-KO and NT control cells with TB40/E and measured the protein levels of ELOVL7 and ELOVL5 at 72 and 96 hpi. In HCMV-infected PERK-KO cells, the levels of ELOVL7 were >2-fold lower than in infected NT cells (Fig. 11F-G). However, the levels of ELOVL5 in HCMV-infected PERK-KO and NT cells were similar (Fig. 11H-I). We conclude that PERK is necessary for HCMV infection-related increase in ELOVL7, but not ELOVL5, protein levels.

**Figure 11.**
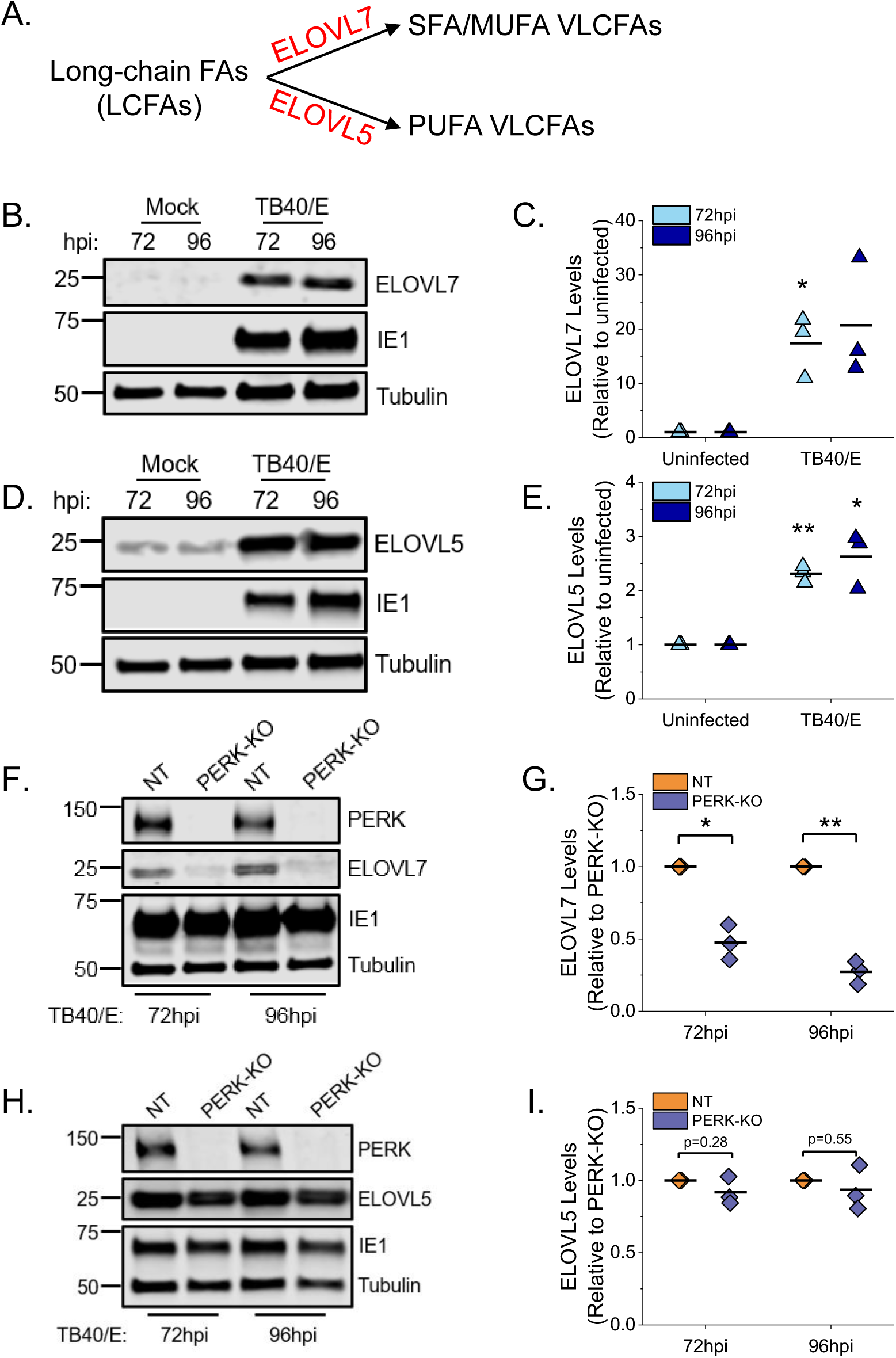
PERK enhances protein levels of ELOVL7 but not ELOVL5. **(A)** Schematic showing ELOVL7 and ELOVL5 elongation of SFA/MUFA VLCFAs and PUFA VLCFAs, respectively. **(B-C)** Western blot analysis and protein quantification for ELOVL7 in HCMV-infected and uninfected HFF cells. ELOVL7 protein levels were normalized to tubulin. **(D-E)** Western blot analysis and protein quantification for ELOVL5 in HCMV-infected and uninfected HFF cells. **(F-G)** Western blot analysis and protein quantification for ELOVL7 in TB40/E-infected PERK-KO-c1 and NT cells. **(H-I)** Western blot analysis and protein quantification for ELOVL5 in TB40/E-infected PERK-KO-c1 and NT cells. *p<0.05, **p<0.01, one-sample t test. MOI 1. N=3

## DISCUSSION

Metabolic reprogramming by HCMV requires several host factors, including those associated with cellular stress responses (4, 28–31). While both HCMV infection and ER-stress are known to regulate lipid metabolism (3, 7, 8, 10, 12, 13, 32, 33), the role of ER-stress in HCMV reprogramming of lipid synthesis is poorly defined. The increase in lipid metabolism following HCMV infection depends on several host factors, including the ER-stress related kinase, PERK (7). However, the effects of PERK on the metabolism of specific classes or types of lipids are unknown. In this study, we used CRISPR/Cas9 engineering, virus replication assays, and lipidomics to define how PERK regulates lipid levels in HCMV-infected cells.

We find that PERK protein levels and activity, as measured by ATF4 protein levels, are increased by 48 hpi (Fig. 1). Many of the mechanisms used by HCMV to induce host stress responses, including PERK, are unknown. The viral ER-resident glycoprotein pUL148 activates PERK (5) and reorganizes the ER membrane (34). However, infection with AD169, which lacks the *UL148* gene, promotes PERK, indicating that HCMV encodes multiple mechanisms to activate PERK (8). Another HCMV protein pUL37x1 induces the release of Ca^2+^ from the ER into the cytosol (35), which may potentially cause ER stress (3, 36). We found that pUL37x1 promotes PERK and ELOVL7 protein levels (37). PL-VLCFAs with SFA/MUFA tails are reduced in cells infected with a mutant virus that lacks the *UL37x1* gene (8), further suggesting that Ca^2+^ flux may be a mechanism involved in inducing ER stress and remodeling of the host lipidome following HCMV infection. Another viral protein, pUL38, that activates PERK (22) also promotes ELOVL7 expression (13), providing further evidence of a connection between PERK and FA elongation. These observations suggest that HCMV encodes several mechanisms to activate ER-stress, each of which may contribute to the reprogramming of lipid metabolism following infection.

HCMV infection increases the flow of carbons from nutrients to lipid synthesis. Carbons from glucose and acetate are used to generate FAs, including VLCFAs made by ELOVLs (8, 12, 13, 26). Using AD169, we previously demonstrated that HCMV infection increases ELOVL5 and ELOVL7 transcripts, ELOVL7 protein levels, FA elongation and VLCFA synthesis, the abundance of PLs with SFA/MUFA tails, and the levels of PERK protein (8, 13, 14). The current study confirms that HCMV infection increases FA elongation, VLCFA synthesis, the abundance of PLs with SFA/MUFA VLCFA tails, and PERK protein level and activity using the more-clinically relevant TB40/E strain. Moreover, we further find that HCMV infection increases the cellular abundance of most DGs and TGs with SFA/MUFA VLCFA tails (Fig. 3A-C). Using CRISPR/Cas9 engineered PERK-KO cells, we demonstrate that PERK contributes to the regulation of lipids levels in HCMV-infected cells. In PERK-KO cells, we observe a rise in the relative levels of DGs, TGs, and PLs with PUFA tails following HCMV infection (Fig. 3F-H, 4, 5, 7, 8D-E, 10). PUFA tails are elongated by ELOVL5 (15, 19). HCMV-infected cells have a higher level of ELOVL5 protein than uninfected cells (Fig. 5D-E). Consistent with an increase in lipids with PUFA tails in infected PERK-KO cells, we observe an increase in ELOVL5 protein levels following HCMV infection independent of PERK (Fig. 5H-I). In HCMV-infected PERK-KO cells, the levels of PLs with SFA/MUFA VLCFA tails are reduced relative to infected control cells (Figs. 8D-E and 9A-B), suggesting that PERK promotes the synthesis of PLs with SFA/MUFA tails. SFA/MUFA VLCFAs are synthesized by ELOVL7 (13, 15, 16, 27). ELOVL7 promotes HCMV replication by supporting the release of infectious virions (13). In this study, we find that PERK is necessary for efficient replication of HCMV (Fig. 2C-D and S1D-E), at least in part, by promoting the infectivity of released virus particles (Fig. 2E). Moreover, we show that PERK helps promote ELOVL7 protein and PLs with SFA/MUFA VLCFA tails levels in HCMV-infected cells (Fig. 8-11). Further, the loss of PERK reduces the particle-to-infectious virion ratio (Fig. 2E), similar to that previously observed when ELOVL7 is depleted (13). Our observations collectively show that the regulation of lipid metabolism in HCMV infected cells involves a PERK-dependent mechanism.

In HCMV-infected PERK-KO cells, ELOVL7 levels are significantly reduced, but not ELOVL5 protein levels (Fig. 11). This loss in the balance of ELOVL5 and ELOVL7 proteins likely reduces the available pool of SFA/MUFA VLCFAs relative to the pool of PUFA VLCFAs. This shift in the saturation status in the pool of available FA tails could explain why lipids with PUFA tails increase in PERK-KO cells (Fig. 3D-H). Based on our observations, we propose a model wherein PERK, through ELOVL7 activity, promotes the synthesis of SFA/MUFA VLCFAs to balance the elongation of PUFAs made by ELOVL5. In HCMV-infected cells, ELOVL5 generates a pool of PUFAs that can be used to synthesize lipids, including DGs, TGs, and PLs (Fig. 12). To ensure the synthesis of lipids with SFA/MUFA VLCFA tails, PERK increases ELOVL7 activity and the generation of SFA/MUFA tails. We further propose that the synthesis of PLs with SFA/MUFA VLCFA tails supports the release of infectious virions. This idea is further supported by the observation that the virus envelope of HCMV is enriched with PLs, with SFA/MUFA VLCFA tails (13, 38). While HCMV replication depends on ELOVL7 activity, the overproduction of saturated VLCFAs by ELOVL7 is cytotoxic (39) and PC, DG, and TG lipids with SFA/MUFA VLCFA tails increase during necroptosis (40). An increase in lipotoxicity could reduce viral replication if cell death occurs before the production of viral progeny. To overcome cell death associated with viral infection, HCMV encodes several inhibitors of cell death pathways (41, 42). The relationship between ER stress and lipid metabolism is bidirectional: ER stress regulates lipid metabolism and lipids can induce ER stress. An overabundance of SFAs induces ER stress, which can be mollified by PUFAs (43–46). It is possible that HCMV-infected cells maintain a balance between PUFAs and SFA/MUFAs to limit lipotoxic stress. If this is the case, then balancing the lipid products of ELOVL5 and ELOVL7 would support HCMV replication. Currently, the importance of ELOVL5 and lipids with PUFA tails in HCMV replication is unknown and is of interest for further study.

**Figure 12.**
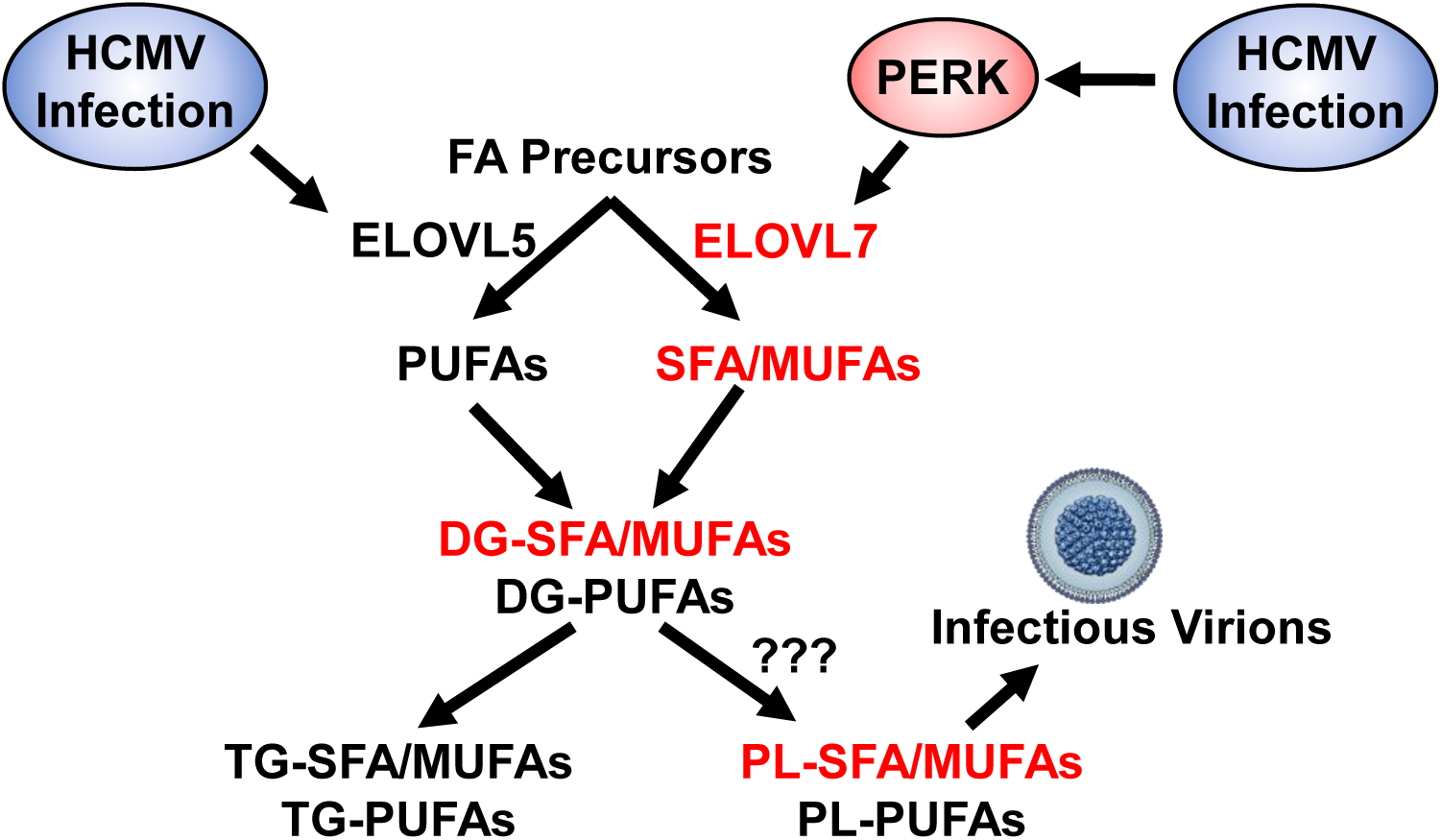
Model for PERK balancing of lipids with SFA/MUFA and PUFA tails through regulation of ELOVL7. HCMV infection increases the elongation of FAs by promoting ELOVL5 and ELOVL7. In HCMV-infected cells, PERK activity promotes ELOVL7, increasing the synthesis of SFAs and MUFAs that are used to generate DGs with SFA/MUFA tails. PERK may further promote lipid synthesis by increasing the flow of DGs to PLs. Further, PLs with SFA/MUFA tails are used to generate infectious virions. In this way, PERK—and more broadly, ER-stress—may help generate membranes necessary for HCMV replication.

We find that, in general, PERK supports the increase in the levels of PLs with SFA/MUFA VLCFA tails following HCMV infection, but not necessarily TGs with SFA/MUFA VLCFA (Fig. 9 compared to Fig. 6). This observatoin could indicate that in addition to regulating ELOVLs, PERK may also promote the flow of DGs to PLs rather than TGs. PERK may affect the flow of lipids through the synthesis pathway by acting directly on the lipid head group. PERK has been reported to phosphorylate DG to generate phosphatidic acid (PA) (23). Since PA lipids are upstream of DGs in the TGs and PLs synthesis pathways, DG conversion to PA by PERK would reduce the synthesis of TGs and PLs. Since the levels of TGs and PLs were increased in PERK-KO cells, it seems unlikely that PERK is phosphorylating DGs during HCMV replication. Nevertheless, at this stage, we cannot rule out if PERK can promote the flow of DGs to PLs, either directly or indirectly.

PERK can regulate FA synthesis by inducing the maturation of sterol regulatory-element binding proteins (SREBPs) (7). SREBPs are transcription factors that promote the expression of lipogenic genes, including ELOVLs. However, overexpression of SREBP1 in PERK-knockdown cells only partially rescued the loss in lipid synthesis, suggesting that PERK uses additional mechanisms to regulate lipid metabolism (7). Moreover, PERK promotes the transcriptional activity of ATF4, a transcription factor that controls amino acid and lipid metabolism during stress (47–52). In white adipose tissue in mice, ATF4 upregulates the expression of FA synthase (FAS) and stearoyl-CoA desaturase-1 (SCD1) (50). In HCMV-infected cells, PERK activity increases ATF4 protein levels (Fig. 2) (5, 21, 22), allowing for a possible role of ATF4 in HCMV-induced lipogenesis. If ATF4 is acting downstream of PERK to regulate lipid metabolism, it may contribute to the balancing SFAs/MUFAs and PUFAs. While continued work is necessary to further define the mechanistic details in PERK regulation of lipid metabolism, our findings demonstrate a previously unknown role of PERK differentially regulating the activity of ELOVLs.

Humans encode seven ELOVLs (ELOVL1-7). While gene expression of all ELOVLs is known to be differentially regulated in a tissue and cell type-dependent manner (15), the mechanisms that differentially regulate them are undefined. Our observation that ELOVL5 and ELOVL7 protein levels are differentially regulated by a PERK-dependent mechanism (Fig. 11) provides insight into cellular control of ELOVLs. Importantly, the observations reported in this study suggest that the differential control of ELOVLs is regulated by a stress response induced by viral infection. It may be possible that stress induced by non-viral or non-infectious triggers will similarly lead to the differential regulation of ELOVLs. In this case, our findings would provide additional evidence supporting the idea that HCMV induces stress to promote virus replication (4). It is noteworthy that the loss of PERK had little to no effect on the levels of lipids in uninfected cells (Fig. 3D-E and Fig. 8C). It is possible that this is due to a lack of stress stimuli in our uninfected cells, and PERK-dependent lipidome changes may occur in uninfected cells following activation of ER stress. Outside of the context of infection, ELOVL7 is expressed in many tissues with the highest levels observed in the pancreas and prostate (15, 16), and its overexpression is associated with prostate cancer (16). ELOVL7 is also expressed in the brain, and a decrease in its expression correlates with Parkinson’s disease (53). It is unknown if a PERK-dependent mechanism controls ELOVL7 protein levels in these diseases. Nonetheless, our finding that PERK differentially regulates ELOVLs to promote ELOVL7 highlights the need for continued studies to understand the role of ER stress in regulating lipid metabolism in viral and non-viral diseases.

While the complex interactions between ER stress and lipid metabolism remain incompletely understood, the conclusions provided in this study highlight that HCMV replication relies on the regulation of lipid metabolism by a stress response to promote FA elongation and to control the relative levels of lipids in host cells. Following HCMV infection, PERK regulates lipid metabolism by promoting ELOVL7, but not ELOVL5. ELOVL7 supports the infectivity of released HCMV virions, and its SFA VLCFA products are present in the HCMV envelope (13). By regulating ELOVL7 and lipid metabolism, PERK—and more broadly, ER stress—may be necessary for membrane biogenesis needed for HCMV replication. Overall, this study furthers our understanding of the mechanisms by which HCMV induces lipogenesis to ensure virus replication by demonstrating that PERK is an essential factor in HCMV-host metabolism interactions.

## MATERIALS AND METHODS

### Cells, Viruses, and Experimental setup

Human foreskin fibroblast (HFF) cells were cultured in Dulbecco’s modified Eagle’s medium (DMEM) containing 10% fetal bovine serum (FBS), 10 mM HEPES, and penicillin-streptomycin. Prior to infection, cells were maintained at full confluence for 3 days in serum-containing growth medium. Cells were switched to serum-free medium (DMEM, HEPES, and penicillin-streptomycin) the day before infection. Cells were infected with either HCMV AD169 or TB40/E strains. Infections were performed at a multiplicity of infection (MOI) of 1 or 3 infectious unit per cell, depending on the experiment. Fresh media was applied to cells at 48 hpi to maintain sufficient nutrients. All virus stocks expressing green fluorescent protein (GFP) were made by pelleting virions from the supernatant of infected cells through 20% sorbitol. Viruses were resuspended in serum-free DMEM and stored at −80°C. Infectious virus titers were measured as the 50% tissue culture infectious dose (TCID_50_) by visualizing GFP signal using immunofluorescence microscopy or after immunofluorescence detection of immediate-early antigen (IE1). Particle-to-infectious unit ratio was determined by the calculating the ratio between viral DNA and infectious virus yield in cell free supernatants. Determination of virus yields from cell free supernatants is described elsewhere (54). Viral DNA in supernatants was quantified by qPCR as previously described (55).

### Focus Expansion assay measuring cell-to-cell spread of HCMV infection

Cell-to-cell spread was measured by immunofluorescence detection under microscopy as previously described (56, 57). Briefly, 150,000 cells per well of the PERK-KO or NT control cells were seeded in a 12-well plate and grown to full confluency for 3 days. Cells were infected with TB40/E HCMV for 24h. At 24 hpi, cells were overlayed with growth media with methylcellulose for another 8 days. At 5 dpi, fresh media with methylcellulose were added to cells. At 9 dpi, cells were fixed with 4% paraformaldehyde (PFA) for 10min at 4°C. HCMV infected cells were stained with mouse monoclonal anti-IE1 (clone 63-27) and mouse monoclonal anti-pp65 (clone 28-277) and detected by immunofluorescence microscopy. The images were acquired by Axio-Observer.Z1 fluorescence microscope and the Axiovision 4.8 software.

### Generation of PERK Knockout Cells Using CRISPR/Cas9 Engineering

sgRNA sequence specific for human gene PERK was cloned into LentiCRISPR-v2 (58, 59), which is a lentiviral vector co-expressing a mammalian codon-optimized Cas9 nuclease with a sgRNA (26). A sgRNA that does not target any human or HCMV sequence is used for a non-targeting (NT) control. The gRNA sequences used in CRISPR knockout experiments are: PERK-KO c1 (G6), CACCTCAGCGACGCGAGTAC; PERK-KO c2 (E4), TGGAGCGCGCCATCAGCCCG; NT, CGCTTCCGCGGCCCGTTCAA. Lentivirus expressing Cas9 and sgRNA were produced in 293FT cells and transduced into life-extended primary human fibroblasts (HFF-hTERT) followed by drug selection using 2 µg/ml puromycin. Single cell clones were obtained by dilution into a 96-well plate by seeding 1.5 transduced cells and 200 non-transduced cells, as previously described (26). 2 µg/ml puromycin was used for selection once the plate reached 85-90% confluence. PERK CRISPR knockout effect was confirmed by Western blot and sequencing.

### Lipidomics analysis

The abundance of all lipids was measured using liquid chromatography–high-resolution tandem mass spectrometry (LC-MS/MS) as previously described (8). Briefly, cells were washed with PBS and lysed in cold 50% methanol. Lipids were extracted twice by using chloroform and dried under nitrogen gas. Lipids then were resuspended in 100 µl of a 1:1:1 solution of methanol-chloroform-isopropanol per 100,000 cells. For each sample, a total of three wells were used for analysis. Two wells were used for lipid extraction, and one well was used to determine the total number of cells. Samples were normalized according to the total number of live cells at the time of lipid extraction. A “no-cell” control was included for any contaminants during lipid extraction process and were removed from the analysis. During data collection, the samples were stored at 4°C in an autosampler. Buffers or any parameter settings for running samples through LC-MS/MS were used as previously described (8). All data were analyzed using EI-MAVEN and Xcalibur (Thermo Scientific).

### Protein Analysis

Proteins were examined by Western blotting using SDS-PAGE performed with tris-glycine-SDS running buffer. Proteins were seperated using Mini-Protean TGX anyKD or 4 to 20% gels (Bio-Rad) and transferred to an Odyssey nitrocellulose membrane (Li-Cor). Membranes were blocked using 5% milk in tris-buffered saline with 0.05% Tween 20 (TBS-T) and incubated with primary antibodies in the presences of 1% milk TBS-T solution, except for anti-ATF4 and anti-pUL123 which were incubated in the presences of 3% or 5% bovine serum albumin (BSA) in TBS-T, respectively. The following antibodies were used: mouse monoclonal anti-pUL123 (IE1, 1:50 dilution), rabbit monoclonal anti-PERK (Cell Signaling; #3192; 1:600 dilution), monoclonal anti-ATF4 D4B8 (Cell Signaling; #11815; 1:600 dilution), rabbit polyclonal anti-ELOVL7 (Sigma-Aldrich; #SAB3500390; 1:1,000 dilution), rabbit polyclonal anti-ELOVL5 (Sigma-Aldrich; #SAB4502642; 1:500 dilution), rabbit polyclonal anti-β-actin (Proteintech; #20536-1-AP; 1:2,000 dilution), and mouse monoclonal anti-α-tubulin (Sigma-Aldrich; #T6199; 1:2,000 dilution). Blots with mouse monoclonal anti-HCMV, anti-actin, and anti-tubulin antibodies were incubated for 1h at room temperature. All others were incubated overnight at 4°C. Quantification of Western blots was performed using a Li-Cor Odyssey CLx imaging system.

## ACKNOWLEDGEMENTS

We thank James Alwine for helpful feedback related to this project and critical reading of the paper. We thank Lisa Wise for helpful feedback related to the lipidomic and knockout studies and Debbie Mustacich for assisting with CRISPR/Cas9 engineering. This project was supported by startup funds from the University of Arizona Health Sciences, College of Medicine-Tucson, and BIO5 Institute to J.G.P. Additional support for this work was provided by a New Investigator Award (ADHS18-198868) to J.G.P. from the Arizona Biomedical Research Commission, made available through the Arizona Department of Health Services.

## Supplemental Figure Legends

**Figure S1.**
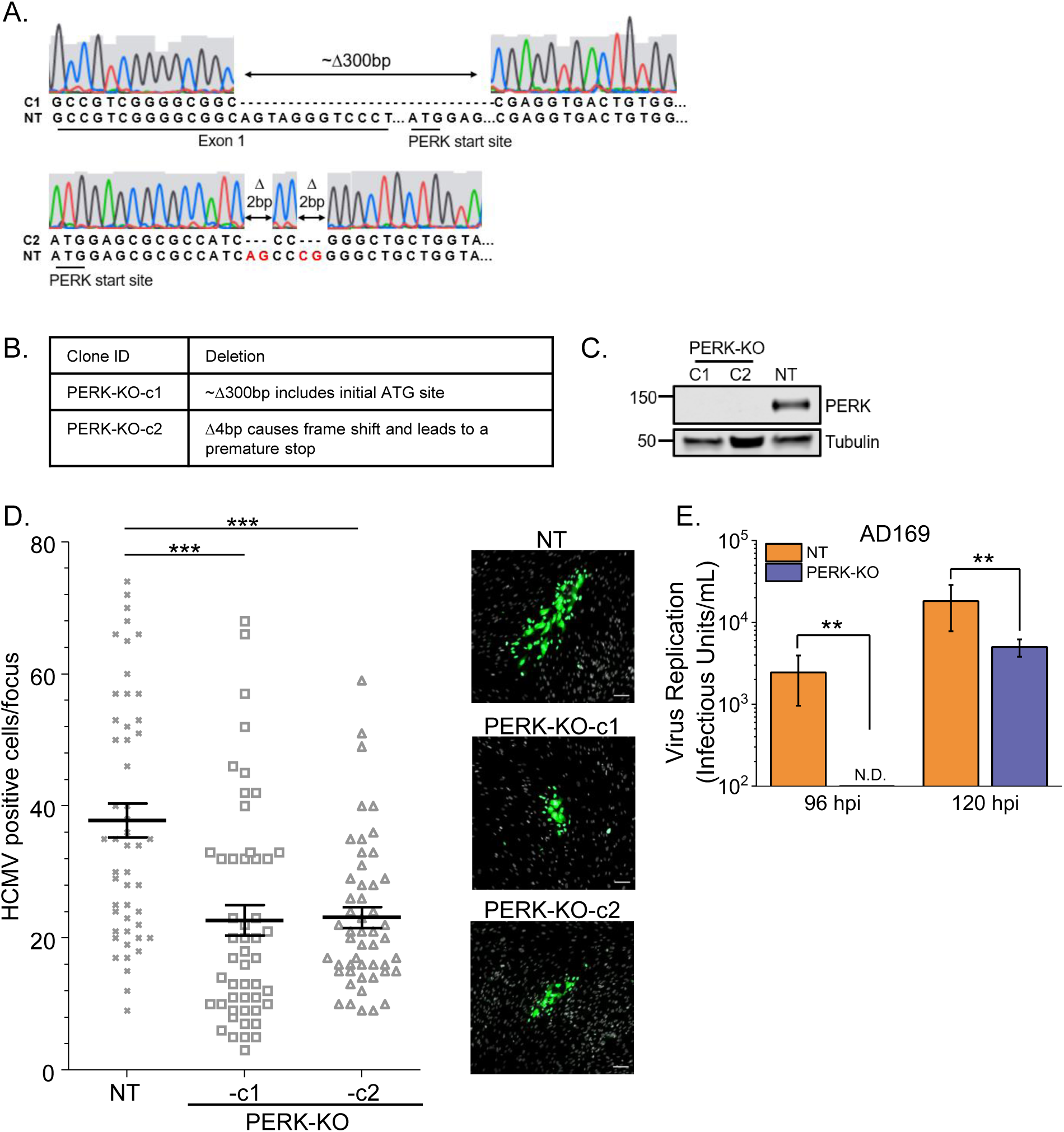
CRISPR/Cas9 knockout of PERK reduces HCMV replication. **(A)** Sequencing results for PERK-KO clone 1 and 2. **(B)** Genotype summary of PERK-KO clones 1 and 2. **(C)** Western analysis of PERK protein levels in PERK-KO clones and control cells that contain a CRISPR/Cas9 non-targeting (NT) gRNA. **(D)** Focus expansion assay measuring TB40/E spread in PERK-KO and NT cells under methylcellulose overlay. Each data point represents the number of HCMV positive cells per focus at 9 days post infection (dpi). Shown are the mean focus sizes and standard deviations of 50 foci for each cell line. Significance testing was performed by a Kruskal-Wallis test followed by a Dunn’s multiple-comparison test (P<0.05). Images of representative HCMV foci in the indicated cell lines after indirect immunofluorescence staining for IE1-antigen and pp65 (green). Cell nuclei were stained with DAPI (grey). The scale bar corresponds to 100 µm. ***p<0.0001. **(E)** At 96 and 120 hpi, AD169 replication in PERK-KO-c2 cells and NT cells infected at MOI 1 was measured by TCID50. Not determined (N.D.) viral titer was below limit of detection. Two-sample t test, **p<0.01.

**Figure S2.**
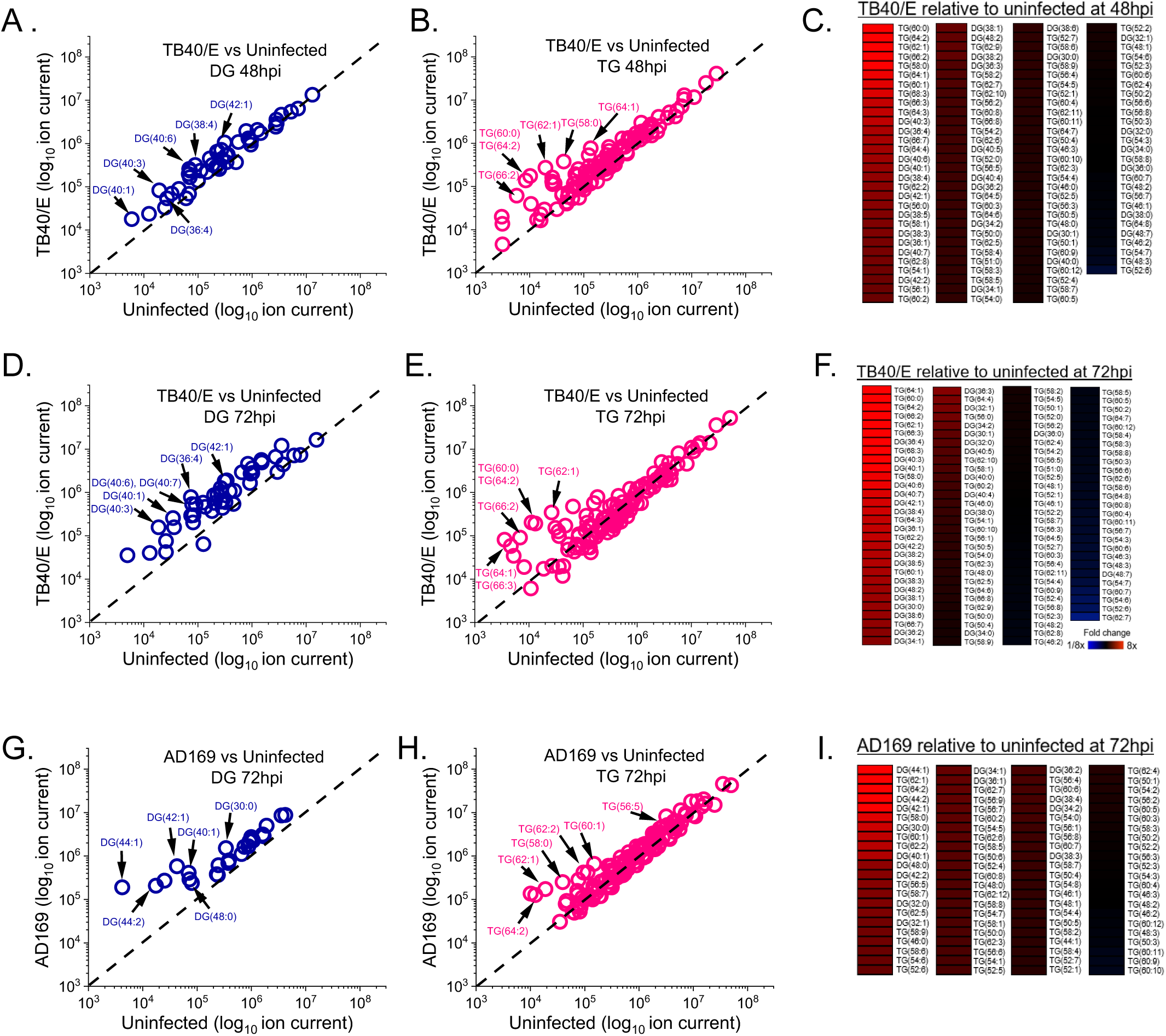
HCMV infection increases DG and TG levels. **(A)** Relative levels of DGs in TB40/E-infected cells and uninfected cells at 48 hpi. Sodiated and ammoniated adducts of DGs were measured by LC-MS/MS following normalization by cell number. Each dot in the plot represents the level of an adduct form of a DG lipid in TB40/E-infected cells relative to its level in uninfected cells. The dash line represents a relative level of 1 (e.g., the level in infected cells is equal to the level in uninfected cells). **(B)** Relative levels of TGs in TB40/E-infected cells and uninfected cells at 48 hpi. TG data was analyzed and visualized using the same methods described in part A. **(C)** Changes in the relative levels of DGs and TGs in HCMV-infected cells and uninfected cells were quantified by averaging the relative levels of the sodiated and ammoniated adducts if both were measured. The averaged relative fold changes were log transformed and visualized as a heatmap. **(D-F)** Relative levels of DGs and TGs in TB40/E-infected cells and uninfected cells at 72 hpi. **(G-I)** The same analysis was performed to determine the relative levels of DGs and TGs in AD169-infected cells and uninfected cells at 72 hpi. MOI=3. N=3.

**Figure S3.**
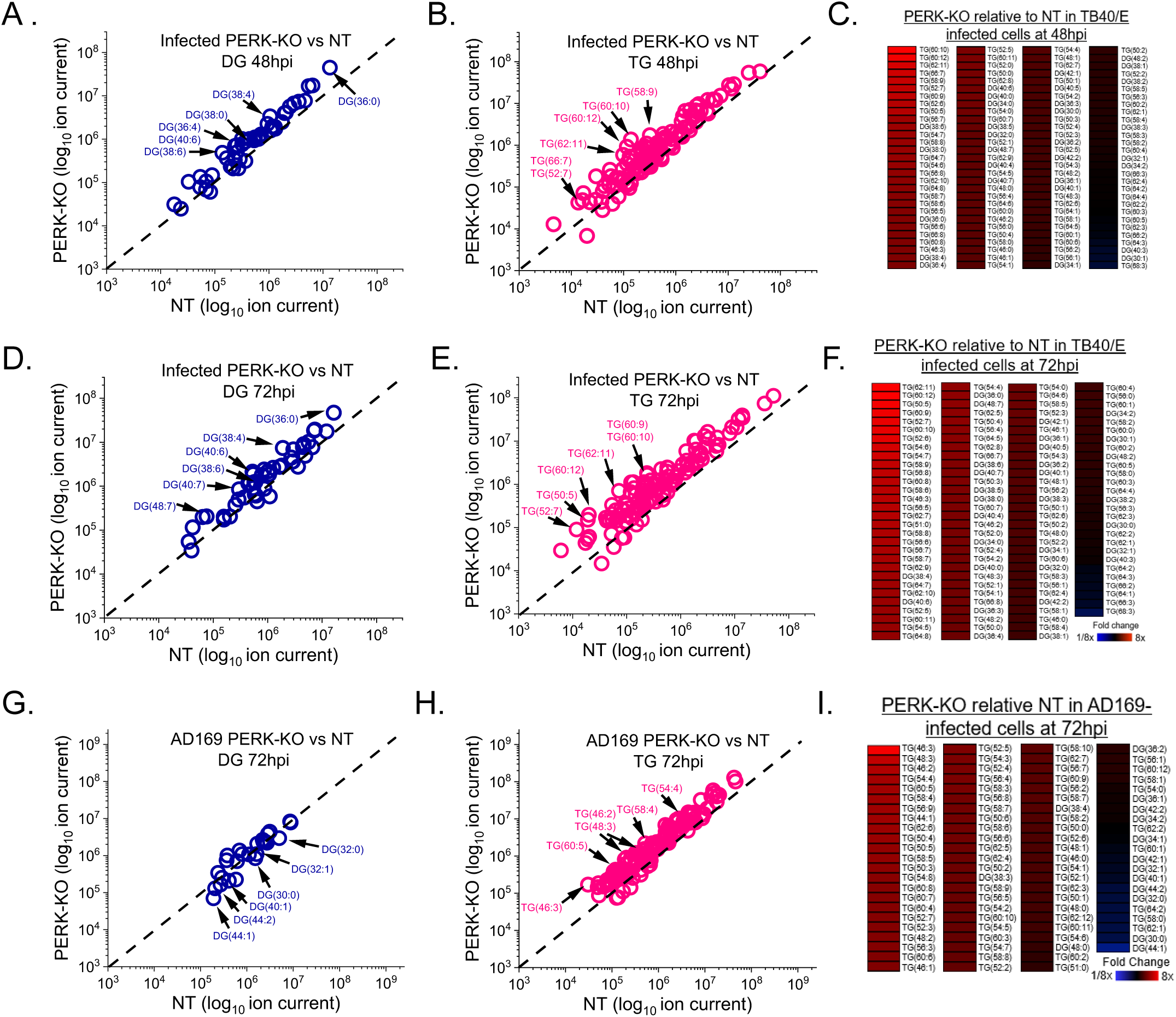
Relative DG and TG lipid levels in HCMV-infected PERK-KO and NT cells. **(A-C)** Relative levels of DGs and TGs in TB40/E-infected PERK-KO and NT control cells at 48 hpi. **(D-F)** Relative levels of DGs and TGs in TB40/E-infected PERK-KO and NT control cells at 72 hpi. **(G-I)** Relative levels of DGs and TGs in AD169-infected PERK-KO and NT control cells at 72 hpi. MOI=3. N=3.

**Figure S4.**
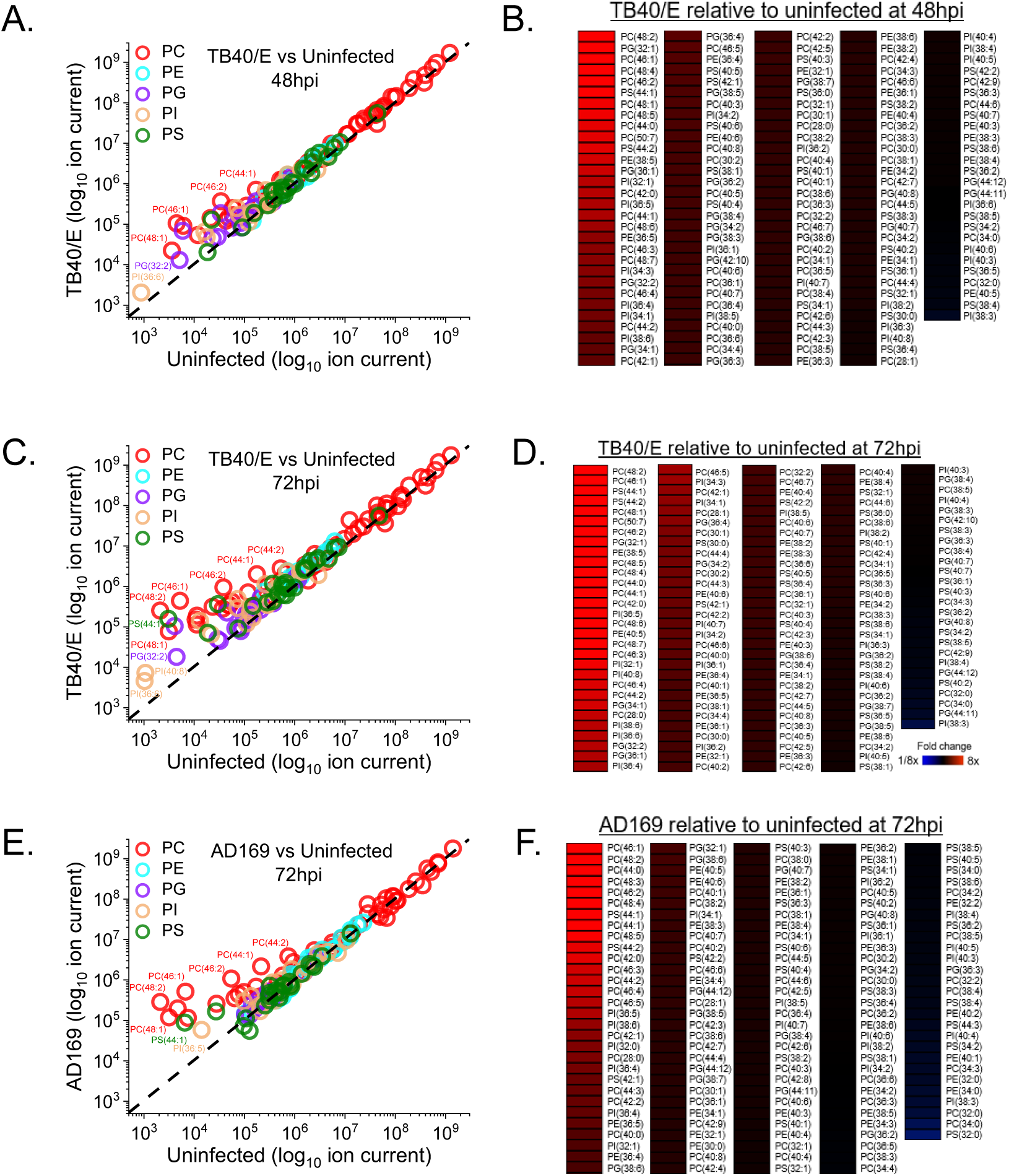
HCMV infection increases PL levels. **(A-B)** Relative levels of PLs in TB40/E-infected and uninfected cells at 48 hpi. **(C-D)** Relative levels of PLs in TB40/E-infected and uninfected cells at 72 hpi. **(E-F)** Relative levels of PLs in AD169-infected and uninfected cells at 72 hpi. MOI=3. N=3.

**Figure S5.**
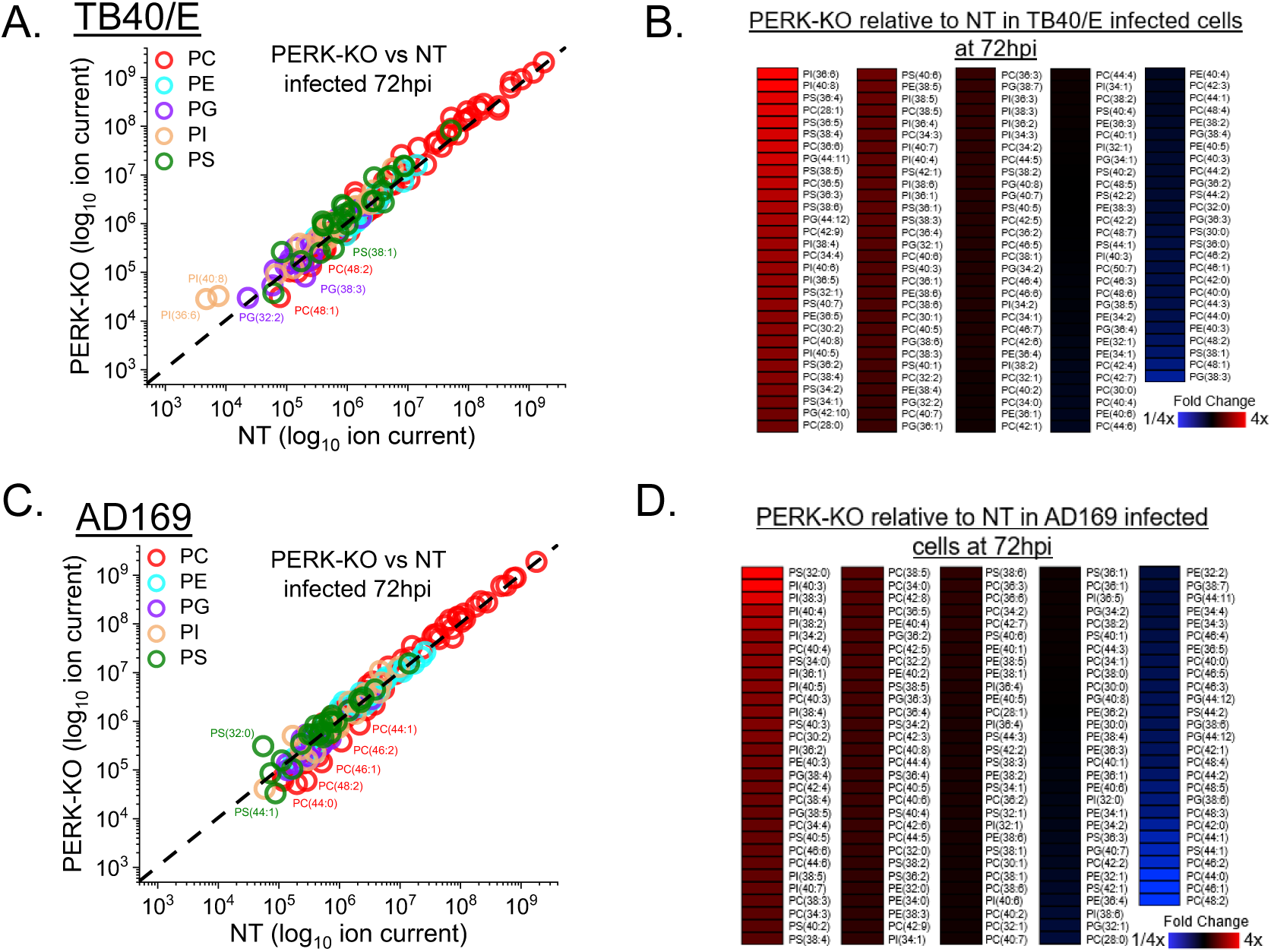
Relative levels of PLs in HCMV-infected PERK-KO and NT cells. **(A-B)** Relative levels of PLs in TB40/E-infected PERK-KO and NT control cells at 72 hpi. **(C-D)** Relative levels of PLs in AD169-infected PERK-KO and NT control cells at 72 hpi. MOI=3. N=3.

